# Regulation of invasion-associated actin dynamics by the *Chlamydia trachomatis* effectors TarP and TmeA

**DOI:** 10.1101/2021.11.04.467231

**Authors:** Matthew D. Romero, Rey A. Carabeo

**Author notes:** Corresponding Author: Department of Pathology and Microbiology, University of Nebraska Medical Center, 985900 Nebraska Medical Center, Omaha, NE.

## Abstract

The obligate intracellular pathogen *Chlamydia trachomatis* manipulates the host actin cytoskeleton to assemble actin-rich structures that drive pathogen entry. The recent discovery of TmeA, which like TarP is an invasion-associated type III effector implicated in actin remodeling, raised questions regarding the nature of their functional interaction. Quantitative live-cell imaging of actin remodeling at invasion sites revealed differences in recruitment and turnover kinetics associated with TarP and TmeA pathways, with the former accounting for most of the robust actin dynamics at invasion sites. TarP-mediated recruitment of the actin nucleators formin and the Arp2/3 complex were crucial for rapid actin kinetics, generating a collaborative positive feedback loop that enhanced their respective actin-nucleating activities within invasion sites. In contrast, Fmn1 is neither recruited to invasion sites nor collaborates with Arp2/3 within the context of TmeA-associated actin recruitment. While the TarP-Fmn1-Arp2/3 signaling axis is responsible for the majority of actin dynamics, its inhibition had similar effects as deletion of TmeA on invasion efficiency, consistent with the proposed model that TarP and TmeA acting on different stages of the same invasion pathway.

**Summary Statement:** Kinetic analysis of actin recruitment during *C. trachomatis* invasion reveals TarP as the major contributor relative to TmeA, via its ability to facilitate collaboration between actin nucleators Formin 1 and Arp2/3.

## Introduction

Infections caused by the obligate intracellular bacterium *Chlamydia trachomatis* are both the leading cause of preventable blindness and the most prevalent bacterial form of sexually transmitted disease worldwide (Malhotra et al., 2013). *Chlamydiae* are obligate intracellular pathogens that feature a biphasic developmental cycle divided between metabolically inactive elementary bodies and the vegetative reticulate bodies. Survival and replication occur in a membrane-bound vacuole known as an inclusion, and therefore, gaining access to the intracellular environment is paramount (Bastidas et al., 2013). Invading elementary bodies initially form a reversible electrostatic attachment onto heparin sulfate proteoglycans on the host cell surface (Su et al., 1996; Zhang and Stephens, 1992). Shortly after, *Chlamydia* engages and activates a multitude of host cell receptors while simultaneously delivering an array of bacterial effectors by a type III secretion system (Chen et al., 2014; Fields et al., 2003; Gitsels et al., 2019). Activation of host receptors and delivery of bacterial effectors during invasion allow *Chlamydia* to exploit regulatory components of the actin cytoskeleton like Arp2/3, Rho GTPases, and vinculin amongst others (Carabeo et al., 2007; Chen et al., 2014; Fadel and Eley, 2008; Gitsels et al., 2019; Hower et al., 2009; Thwaites et al., 2015). By manipulating cytoskeletal regulators, *Chlamydia* induces the formation of actin-rich microstructures on the cell surface that facilitate engulfment of the pathogen (Carabeo et al., 2002; Caven and Carabeo, 2019). These structures can adopt a variety of configurations, including phagocytic cups, membrane ruffles, filopodia, and hypertrophic microvilli (Carabeo et al., 2007; Ford et al., 2018). A distinguishing characteristic of structures generated by *Chlamydia* during invasion is their rapid and highly localized assembly (Carabeo et al., 2002). Whereas *S. typhimurium* and *R. parkeri* recruit actin in a diffuse pattern over the span of 5-15 minutes (Francis et al., 1993; Reed et al., 2014; Zhou et al., 1999), actin remodeling during *Chlamydia* invasion is highly localized and recruitment and disassembly completed within two minutes (Carabeo et al., 2004).

Actin polymerization at the invasion site occurs is due in large part to the secretion of the bacterial effectors TarP and TmeA, which jointly coordinate the assembly of a robust actin remodeling network at *Chlamydia* entry sites. TarP and TmeA accomplish this task in part by their shared ability to recruit the actin nucleator Arp2/3 during invasion. Arp2/3 is a complex of proteins which, upon activation, enhances the rate of actin polymerization by nucleating branched actin filaments along a pre-existing “mother” filament (Bailly et al., 1999; Mullins et al., 1998), and the resulting branched network in turn increases the number of sites for Arp2/3 binding and branched nucleation (Higgs and Pollard, 1999). TarP and TmeA activate the Arp2/3 complex via downstream activation of the nucleation promoting factors WAVE2 and N-WASP, respectively (Carabeo et al., 2007; Faris et al., 2020; Keb et al., 2021; Lane et al., 2008). Both nucleation promoting factors have each been shown to be important for *Chlamydia* engulfment, as targeted disruption of either protein attenuated pathogen uptake (Carabeo et al., 2007; Faris et al., 2020; Keb et al., 2021). Furthermore, siRNA knockdown of WAVE2 prevented the recruitment of actin at entry sites, highlighting the importance of TarP-associated Arp2/3 activation and subsequent actin remodeling in *Chlamydia* pathogenesis (Carabeo et al., 2007).

The actin remodeling network generated by *Chlamydia* enable the assembly of invasion-associated structures which are comprised entirely or in part by elongated bundles of filamentous actin. A class of actin nucleators known as formins participate in the assembly of actin-rich structures by facilitating rapid incorporation of actin monomers onto the barbed end of actin filaments (Shemesh and Kozlov, 2007). In turn, these rapidly elongating filaments provide a highly localized protrusive force that gives rise to actin-rich structures such as filopodia and hypertrophic microvilli (Fattouh et al., 2015; Kage et al., 2017; Yang et al., 2007). Broadly, formins can be divided into two subclasses: diaphanous-related formins (DRFs), which must be activated by Rho-GTPases before they can serve as actin nucleators, and non-DRF formins, which do not require activation to participate in actin filament elongation (Baarlink et al., 2010). Once activated, formins dimerize and incorporate monomeric actin onto the barbed end of an actin filament. Formins have been demonstrated to participate in the invasion of bacterial pathogens including *L. monocytogenes, S. typhimurium*, and *B. burgdorferi* (Boddy et al., 2018; Rengarajan et al., 2016; Williams et al., 2018). Likewise, invading *C. trachomatis* is known to interact with hypertrophic microvillar structures that are enriched with filamentous actin (Carabeo et al., 2002), raising the possibility that *Chlamydia* also utilizes formins during invasion. In addition, formin and its homolog, Diaphanous, along with components of the Arp2/3 complex were identified in an RNA interference screen in Drosophila S2 cells as being important host factors in infection (Elwell et al., 2008).

Here, we implicate three formin family members (Fmn1, mDia1, and mDia2) as important elements of *Chlamydia* invasion. We demonstrate that the recruitment of formin occurred via a TarP-dependent and TmeA-independent process. An extensive yeast two-hybrid screen directed against TarP failed to identify any of the three formin members, indicating that TarP-mediated formin recruitment occurs via indirect interaction. Nevertheless, we observed that formin and the Arp2/3 complex were simultaneously recruited during invasion, implying a functional interaction between these actin nucleators. Indeed, the absence of either nucleator drastically reduced *Chlamydia* entry, which correlated with a significant reduction in the rate of actin recruitment. These findings were recapitulated using the pan-formin inhibitor SMIFH2 and the Arp2/3 inhibitor CK666. Notably, we found that recruitment of Arp2/3 was dramatically diminished when Fmn1 is inhibited, and vice versa, indicating that their respective recruitment depends on the actin network formed by the other. This interdependent recruitment underpins enhanced actin recruitment kinetics, revealing a cross-collaborative interaction which establishes a definitive mechanistic link between rapid actin kinetics and efficient *Chlamydia* invasion. Quantification of actin kinetics also revealed the primacy of TarP signaling as a key element of *Chlamydia* actin recruitment dynamics. Specifically, strains which lacked TarP exhibited profound defects in the recruitment kinetics of actin, Formin 1, and Arp2/3, which were less severe in strains lacking TmeA. Collectively, these findings point to TarP facilitating functional collaboration between formins and Arp2/3 to promote the hallmark rapid actin recruitment and turnover kinetics at the site of invasion.

## Results

### TarP accounts for the majority of invasion-associated actin remodeling

Rapid and robust recruitment of actin is a hallmark of *Chlamydia* invasion, which is coordinated in part by the secreted effectors TarP and TmeA via their ability to recruit actin modulatory proteins during invasion. Recently, it was shown that TarP and TmeA synergistically enhance the in-vitro rate of actin polymerization (Keb et al., 2021). Genetic deletion of TarP (ΔTarP) or TmeA (ΔTmeA) attenuates pathogen entry, which was rescued by cis-complementation of *ct456/tarp* (*cis*-TarP) or *ct694/tmea* (*cis*-TmeA). On the basis of these observations, it was proposed that synergy between TmeA and TarP is an important determinant of both robust actin kinetics and efficient invasion. To evaluate this claim, we quantified the rate and intensity of actin recruitment at the entry sites of wild-type, single-knockout (ΔTarP, ΔTmeA), cis-complemented (*cis*-TarP, *cis*-TmeA), or double-knockout (ΔTmeA/ΔTarP) *Chlamydia* strains (Fig. 1, Video S1). Comparisons of actin recruitment between strains were derived by calculating their respective fold recruitment values, which measures the fold increase of fluorescence intensity at entry sites relative to the fluorescence intensity prior to the arrival of elementary bodies. Importantly, background fluorescence signal was obtained from regions immediately outside of regions of *Chlamydia* entry and subtracted from the specific fluorescence intensity obtained within the entry site. This method of background subtraction accounts for local variations in fluorescence intensity, while calculation of fold recruitment values accounts for potential variations in the expression levels of fluorescent protein. Details of this process can be found in the supplementary data (Fig. S1). Furthermore, we found that the recruitment profiles obtained from exogenously expressed GFP-actin closely resembled the recruitment profile of endogenous actin stained with the far-red dye SiR (Fig. S2), validating the use of either method to monitor actin dynamics.

**Figure 1:**
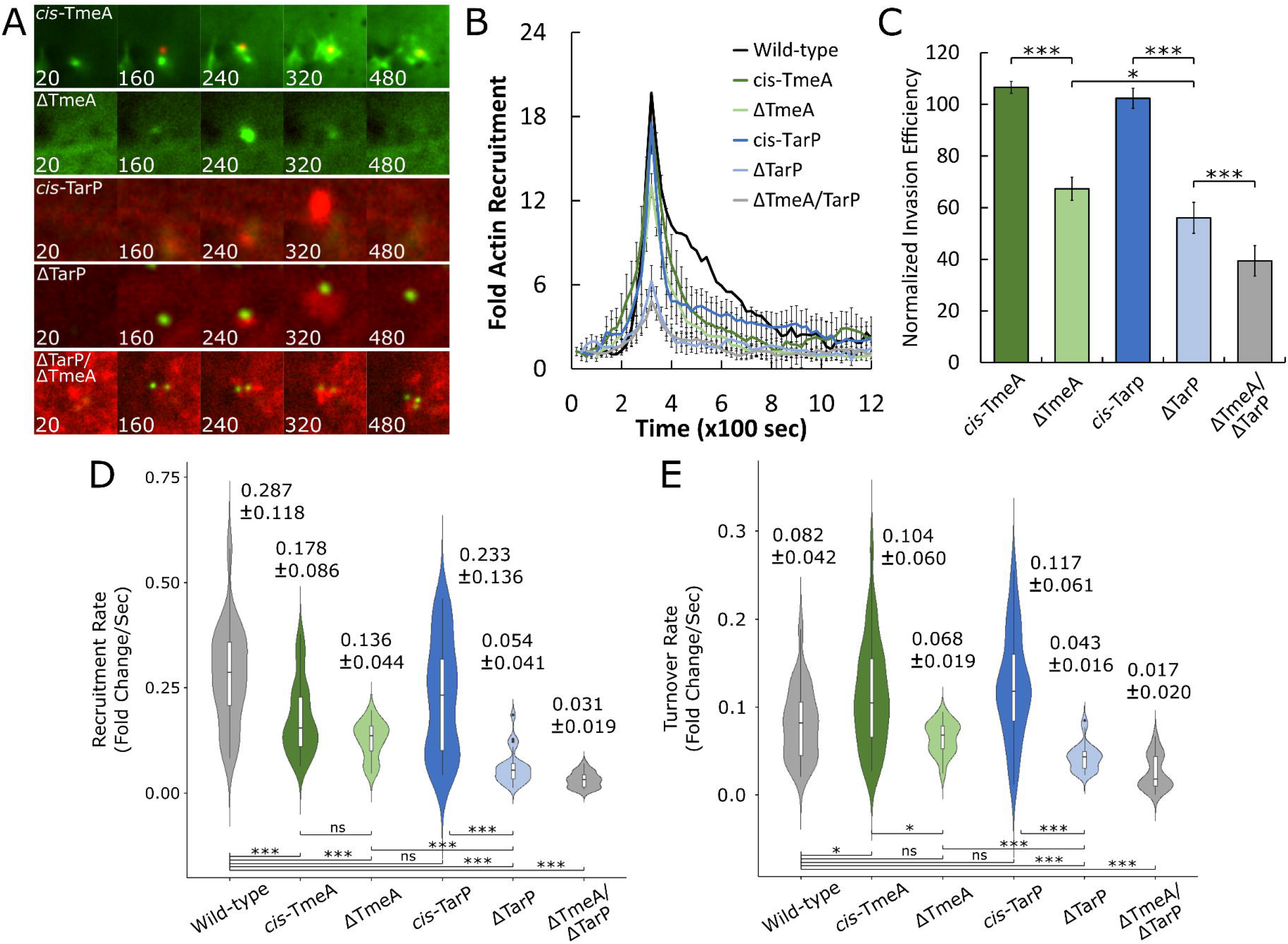
Deletion of TarP and TmeA impair invasion and attenuates kinetics of actin recruitment and turnover. (A,B) Cos7 cells were transfected with Actin-GFP or mRuby-LifeAct for 24 hours prior to infection with wild-type, single-knockout (ΔTarP, ΔTmeA), cis-complemented (*cis*-TarP, *cis*-TmeA), or double-knockout (ΔTmeA/ΔTarP) *Chlamydia* strains at MOI=20. Infection was monitored by live-cell confocal microscopy using a Nikon CSU-W1 spinning disk microscope, obtaining images every 20 seconds for 30 minutes to identify sites exhibiting actin recruitment. (B) Recruitment events were isolated by selecting regions containing invading *Chlamydia* and elevated actin fluorescence. Background fluorescence was subtracted, and fold recruitment was calculated as a function of the fold increase in mean fluorescence intensity of recruited actin compared to basal actin fluorescence. Detailed visualization of this process can be found in Fig. S1. Data is displayed as mean fold recruitment for each timepoint +/-SEM compiled from a minimum N=18 recruitment events. (C) HeLa cells were infected with each *Chlamydia* strain indicated above at MOI=50 and stained using the “in-and-out” method which distinguishes non-internalized EBs from total cell-associated EBs, as described in Materials and Methods. Results were normalized against mean invasion efficiency of wild-type *Chlamydia* and plotted as normalized mean +/- SEM. Data was collected from 15 fields, with each field containing an average of 100 *Chlamydiae*. Statistical significance was determined by T-test. (D,E) All actin recruitment events used to create the averaged plot shown previously (Fig. 1B) were individually divided into recruitment and turnover phases (Fig. S3). Individual rates of (D) recruitment and (E) turnover were plotted on a violin plot with inset boxplot, reporting the median rate +/-SD for each strain tested. Violin plots contain a minimum N=18 individual rates. Statistical significance was determined by Wilcoxon Rank-sum. All data are representative of at least 3 independent experiments, * P ≤ 0.05, ** P ≤ 0.01, *** P ≤ 0.001.

While all strains retained the ability to recruit actin, we noted that the intensity of actin recruitment was diminished upon deletion of TarP and/or TmeA (Fig. 1A,B). Wild-type and cis-complemented strains of *Chlamydia* recruited actin at an average intensity of 18-fold above background, which was reduced by 5-fold following TmeA deletion and 12-fold following TarP deletion. Interestingly, the peak intensity of actin recruitment by the ΔTmeA/ΔTarP double-mutant strain (4.86-fold) was only marginally reduced relative to ΔTarP (6.27-fold), suggesting that TmeA signaling alone is insufficient to generate robust actin kinetics, and that TarP accounts for the majority of the actin signal intensity during invasion. Despite this, and in agreement with earlier studies, we found that deletion of either TmeA or TarP significantly reduced the efficiency of *Chlamydia* internalization (Wild-type=76.2%, ΔTmeA=50.4%, ΔTarP=42.1%), and that deletion of both effectors worsened this defect (ΔTmeA/ΔTarP=29.6%) (Fig. 1C). Collectively, these data indicate that strains lacking TarP and/or TmeA recruit actin suboptimally, which is linked to poor invasion efficiency relative to either wild-type or cis-complemented strains.

In addition to reducing the intensity of actin recruitment, we also noted that the overall actin recruitment profiles of ΔTarP and ΔTmeA strains differed from wild-type *Chlamydia*. In particular, both ΔTarP and ΔTmeA/ΔTarP exhibit atypically transient actin recruitment, while ΔTmeA produces an intermediate profile with slightly lower intensity and duration (Fig. 1B). These observations, coupled with reduced invasion efficiency following TarP or TmeA deletion (Fig. 1C), are further proof of a link between actin kinetics and invasion efficiency. We observed that the recruitment profile of actin exhibited three distinct phases: i) recruitment, in which actin rapidly accumulates, ii) fast turnover, in which actin is rapidly lost from the sites of invasion, likely through depolymerization and iii) slow turnover, in which actin is gradually depolymerized over an extended time period (Fig. 1B). The asynchrony between the time of EB arrival and the start of recruitment required a fixed temporal reference point to enable quantification. For this, we plotted the first derivative values corresponding to the velocity of recruitment and turnover for each timepoint and chose the first timepoint exhibiting positive velocity as the start of recruitment. Further details regarding the criteria used to define recruitment and turnover phases can be found in the supplemental data (Fig. S3).

To compare the kinetics of actin recruitment and turnover between conditions, we employed linear regression to calculate the slope, representing the rate of recruitment and turnover for each invasion event (Fig. S1,S3), and mapped these values onto a violin plot to visualize the distribution of rates for each strain (Fig. 1D,E, Table S1). Deletion of either TmeA or TarP attenuated actin recruitment kinetics by 2-fold and 5-fold, respectively, relative to wild-type *Chlamydia* (ΔTmeA=0.178 fold/sec, ΔTarP=0.054 fold/sec). Actin kinetics were further restricted by simultaneous deletion of both TmeA and TarP (ΔTmeA/ΔTarP=0.031 fold/sec), yielding a nearly 10-fold reduction in the rate of actin recruitment. Altogether, these data suggest that optimal rapid actin recruitment is an effector-driven process that relies on both TarP and TmeA. Furthermore, we observed that actin kinetics were more severely attenuated in strains that lack TarP (ΔTarP, ΔTmeA/ΔTarP), suggesting that TarP is the major determinant of actin recruitment kinetics, and that TmeA has a minor role.

### Formin 1 is involved in TarP-associated actin dynamics

We investigated the basis for the greater impact of TarP on actin recruitment and tested the hypothesis that TarP could be engaging multiple nucleators of actin polymerization. Formin family members, e.g. formin and Diaphanous were identified in an RNA interference screen in Drosophila S2 cells for host factors necessary for infection (Elwell et al., 2008). The need for actin remodeling during invasion raises the possibility that the mammalian equivalents Fmn1, mDia1, and mDia2 are involved in this process. Furthermore, analysis of formin recruitment within the context of the TarP- and TmeA-deficient mutant strains may provide a basis for the greater role of TarP in actin remodeling dynamics.

Since multiple formin species are implicated as invasion-associated factors, we utilized strategies which simultaneously impair their actin nucleating function to mitigate the possibility of compensatory interactions. To this end, we exposed Cos7 cells to the pan-formin inhibitor SMIFH2 (10 μM), which prevents actin monomer binding onto formin species by occupying their conserved formin-homology 2 (FH2) domain, and quantified *Chlamydia* invasion efficiency at 10 min post-infection. At this dose, the reported off-target effects of SMIFH2 at 100 μM on myosin II activity was not observed based on retention of myosin II-dependent focal adhesions and stress fibers in treated mock-infected Cos7 cells (Fig. S4) (Nishimura et al., 2021, 2). Quantitative invasion assay revealed a 39% decrease in invasion efficiency following SMIFH2 treatment relative to mock-treated samples (Fig. 2A). Additionally, simultaneous knockdown of Fmn1, mDia1, and mDia2 using a pooled esiRNA treatment (3x esiRNA) reduced invasion efficiency by 45% compared to both mock- and scramble RNA-treated samples. We note that individual knockdown of each formin species reduced invasion efficiency by approximately 20%, indicating functional compensation between formins (Fig. S5E).

**Figure 2:**
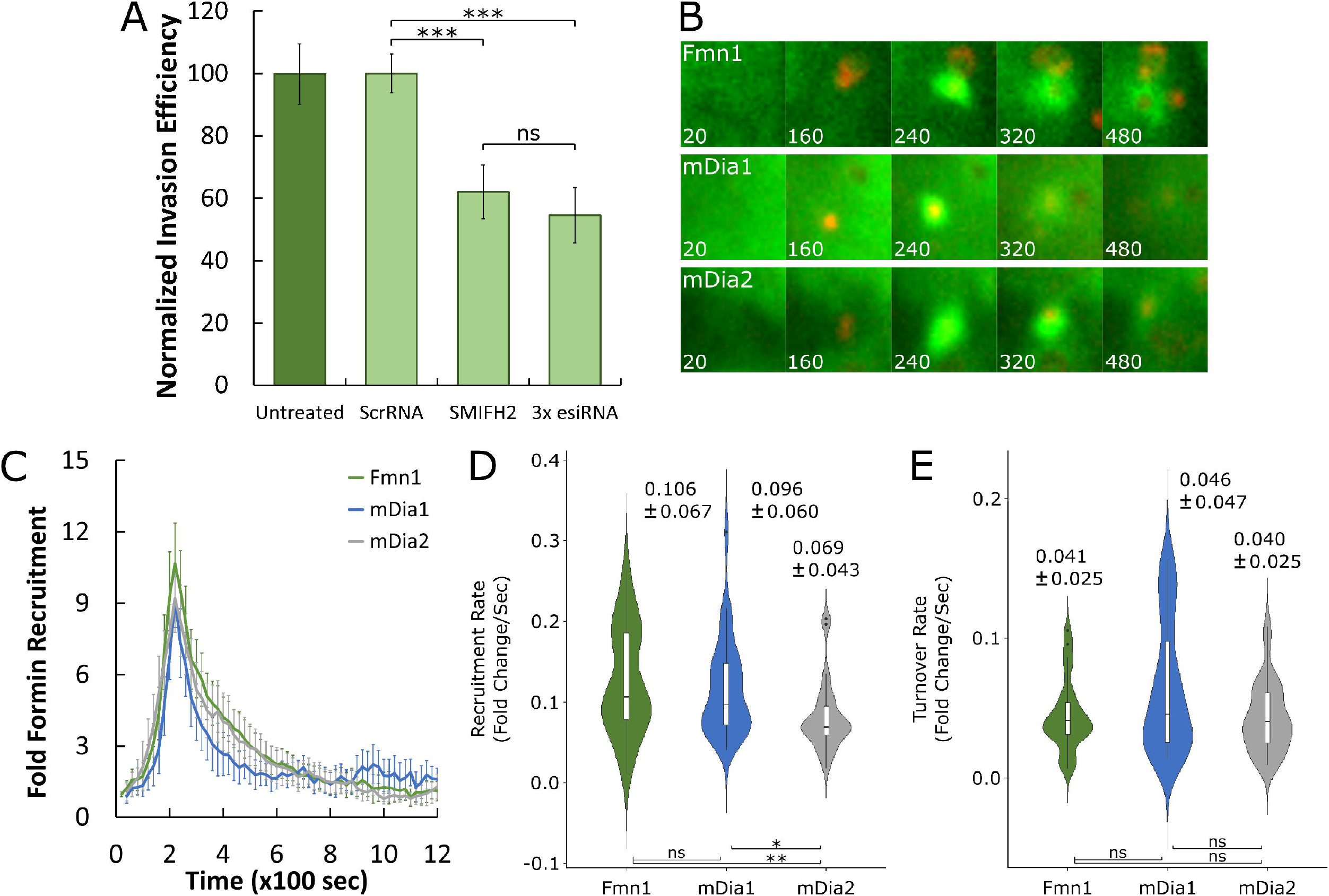
Multiple formin species are recruited during invasion and promote *Chlamydia* internalization. (A) Expression of Fmn1, mDia1, and mDia2 within HeLa cells was simultaneously repressed via RNA interference, using a pooled mixture containing 100ng of three esiRNAs (3x esiRNA) targeting transcripts of each aforementioned gene. Pooled esiRNA or scramble RNA control was transfected into HeLa cells and incubated for 48 hours prior to infection with wild-type *C. trachomatis* elementary bodies at MOI=50. Concurrently, HeLa cells were mock-treated or treated with the pan-formin inhibitor SMIFH2 (10μM) for 1 hour prior to infection (MOI=50). Infected cells were fixed with 4% paraformaldehyde at 10 minutes post infection and stained to distinguish internalized elementary bodies using the method described earlier (Fig. 1C). Results were normalized against mean invasion efficiency of mock-treated *Chlamydia* and plotted as normalized mean +/-SEM. Data was collected from 15 fields, with each field containing an average of 100 *Chlamydiae*. Statistical significance was determined by T-test. (B,C) Cos7 cells were transfected with GFP-Fmn1, mEmerald-mDia1, or mEmerald-mDia2 for 24 hours prior to infection with wild-type *Chlamydia* stained with red-fluorescent dye (CMTPX) at MOI=20. Infection was monitored by live-cell confocal microscopy, obtaining images every 20 seconds for 30 minutes to identify sites exhibiting formin recruitment. Fluorescence intensity of formin recruitment was measured as described earlier (Fig. 1B) and plotted as mean fold recruitment for each timepoint +/-SEM compiled from a minimum N=15 recruitment events. (D,E) Recruitment and turnover kinetics were analyzed for Fmn1, mDia1, and mDia2 recruitment curves using the same methodology described in Fig. 1D,E. Violin plots contain a minimum N=15 individual rates, reporting the median rate +/-SD. Statistical significance was determined by Wilcoxon Rank-sum. All data are representative of at least 3 independent experiments, * P ≤ 0.05, ** p ≤ 0.01, *** P ≤ 0.001.

Given that *Chlamydia* invasion is attenuated by depletion or inhibition of Fmn1, mDia1, and mDia2, we chose to monitor the involvement of these proteins during pathogen entry. Time-lapse images of Cos7 cells transfected to individually express fluorescence-tagged versions of Fmn1, mDia1, and mDia2 were collected and assembled as videos, revealing that each of these proteins was recruited at *Chlamydia* entry sites (Fig. 2B). Moreover, upon plotting their specific fluorescence intensity of over time, we noted that each formin species exhibited rapid, robust, and transient recruitment (Fig. 2C). Using the same method employed to quantify actin kinetics (Fig. 1D,E; Fig. S3), we compared the recruitment and turnover kinetics of recruited formins (Fig. 2D,E). Although the rate of mDia2 recruitment (0.069 fold/sec) was marginally slower than Fmn1 (0.106 fold/sec) and mDia1 (0.096 fold/sec), we report that each formin species possesses largely comparable recruitment kinetics (Fig. 2D). A similar trend persists in their turnover kinetics, as Fmn1 (0.041 fold/sec), mDia1 (0.046 fold/sec) and mDia2 (0.040 fold/sec) each had comparable turnover rates (Fig. 2E). Collectively, these data imply that multiple formin species are recruited during *Chlamydia* invasion, that their recruitment supports pathogen internalization, and that their activities are likely functionally redundant.

Provided the central importance of actin remodeling toward *Chlamydia* invasion, it is highly probable that formins assist in pathogen entry by enhancing actin remodeling kinetics. To evaluate this claim, we measured the effects of SMIFH2 treatment on actin dynamics within entry sites via time-lapse live-cell imaging. Using the same methods described earlier (Fig. S1, S3), we quantified the specific fluorescence intensity of GFP-actin recruited by *Chlamydia* in the presence and absence of SMIFH2 and plotted this data as fold actin recruitment (Fig. 3A). Inhibition of formins yielded a substantial defect to actin recruitment, reducing peak actin intensity by 11-fold (Mock = 19.69-fold vs. SMIFH2 = 8.99-fold). Additionally, the actin recruitment kinetics generated by each experimental group differed significantly in all phases, e.g. recruitment (0.285 fold/sec vs. 0.095 fold/sec), fast turnover (0.108 fold/sec vs. 0.031 fold/sec), and slow turnover (Fig. 3B,C). We were unable to evaluate whether simultaneous knockdown of Fmn1, mDia1 and mDia2 via pooled esiRNA treatment yielded a comparable defect to actin kinetics, as this treatment rendered cells highly susceptible to phototoxicity, and are thus unsuitable for live-cell imaging (Fig. S5D). Nevertheless, these data indicate that the principal cause of reduced invasion efficiency following disruption of formins is due to deficient actin remodeling.

**Figure 3:**
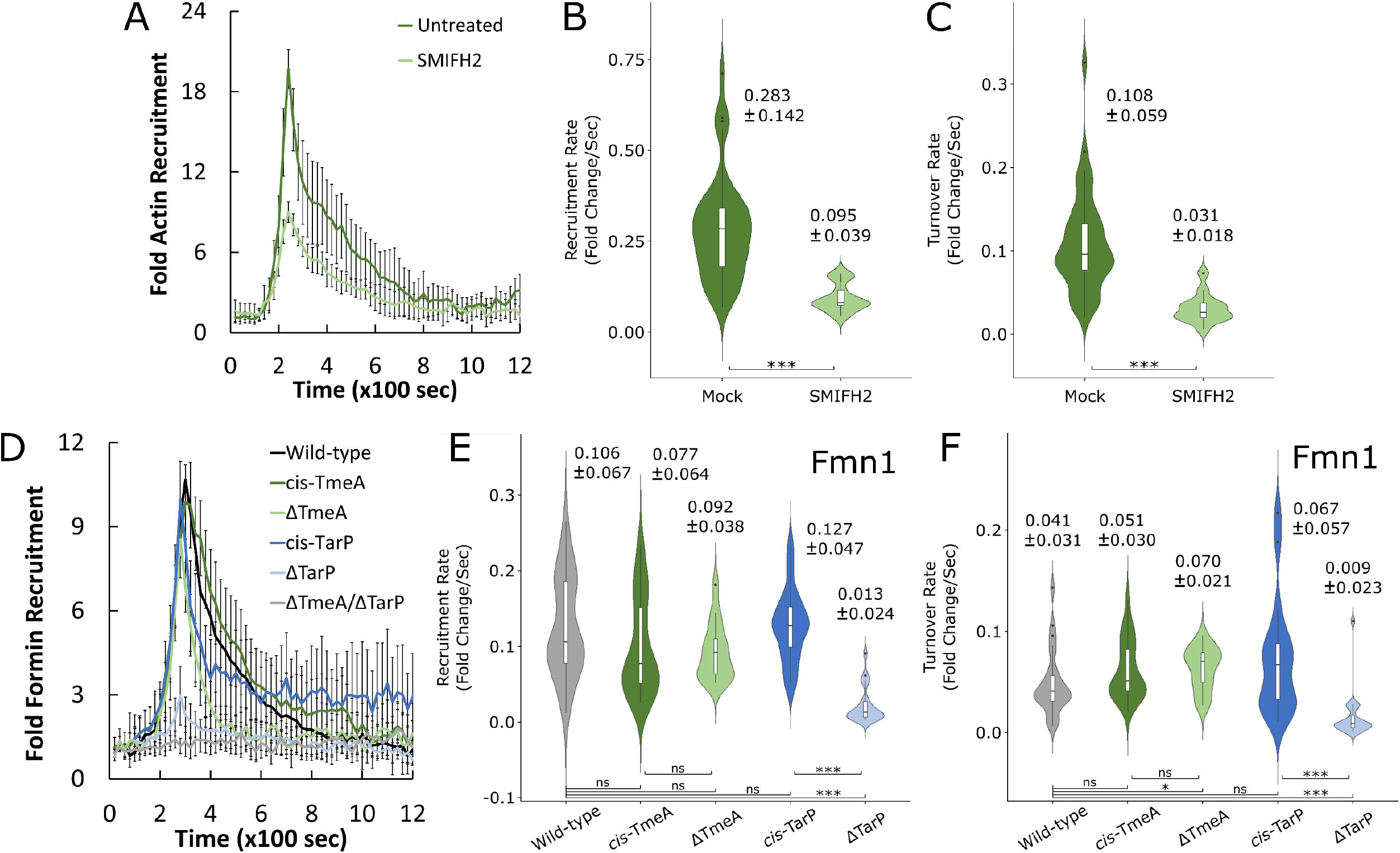
TarP-dependent recruitment of formin is required for rapid actin kinetics. (A,B) Cos7 cells were transfected with Actin-GFP for 24 hours prior to application of the pan-formin inhibitor SMIFH2 (10μM) or mock DMSO control for 1 hour, followed by infection with wild-type CMTPX-stained *Chlamydia* (MOI=20). Infection was monitored by live-cell confocal microscopy, and fluorescence intensity of actin recruitment was measured using methods described earlier (Fig. 1A,B), plotting the mean fold recruitment for each timepoint +/-SEM compiled from a minimum N=16 recruitment events. (B,C) Actin recruitment and turnover kinetics were analyzed for both mock and SMIFH2 treated groups using the same methodology described in Fig. 1D,E. Violin plots contain a minimum N=16 individual rates, reporting the median rate +/-SD. (D) Cos7 cells were transfected with Fmn1-GFP or Fmn1-mCherry for 24 hours prior to infection with the indicated *Chlamydia* strains at MOI=20. Infection was monitored by live-cell confocal microscopy using a Nikon CSU-W1 spinning disk microscope, obtaining images every 20 seconds for 30 minutes to identify sites exhibiting Fmn1 recruitment. Fluorescence intensity of Fmn1 recruitment was measured as described earlier (Fig. 1B) and plotted as mean fold recruitment for each timepoint +/-SEM compiled from a minimum N=17 recruitment events. (E,F) Kinetics of Fmn1 recruitment and turnover were analyzed for each strain indicated above using the same methodology described earlier. Violin plots contain a minimum N=17 individual rates, reporting the median rate +/-SD. Statistical significance was determined by Wilcoxon Rank-sum. All data are representative of at least 3 independent experiments, * P ≤ 0.05, ** P ≤ 0.01, *** P ≤ 0.001.

We observed earlier in this study that strains which lack TarP (ΔTarP, ΔTmeA/ΔTarP) exhibited more significant impediments in both actin recruitment and turnover than either wild-type or TmeA-deficient (ΔTmeA) strains (Fig. 1B,D,E). One possible explanation for this difference may be that formins, which enhance actin remodeling kinetics, are recruited in a TarP-specific manner. As such, we tested whether the recruitment of formin was contingent upon the presence of TarP or TmeA (Fig. 3D-F), focusing our investigation on the recruitment of Fmn1 since this protein was specifically identified in an RNA interference screen of host factors necessary for infection (Elwell et al., 2008). We observed the robust recruitment of Fmn1 at the entry sites of wild-type, cis-complemented, and ΔTmeA strains, which was absent in ΔTarP and ΔTmeA/ΔTarP strains, suggesting that Fmn1 recruitment is TarP-dependent (Fig. 3D). Interestingly, an extensive yeast two-hybrid screen directed against TarP failed to identify Fmn1, mDia1, and mDia2, suggesting that formins do not directly interact with TarP (Table S1). As such, it is likely that TarP-mediated recruitment of formins occurs indirectly, however the precise factors which mediate this interaction remain unknown. TmeA deletion did not appreciably alter the recruitment intensity of Fmn1 relative to either *cis*-TmeA or wild-type *Chlamydia* (wild-type=10.7-fold, *cis*-TmeA=9.9-fold, ΔTmeA=8.5-fold), indicating that recruitment of Fmn1 is mediated primarily by TarP and largely independent of TmeA. In further support of this, we found that Fmn1 recruitment kinetics were not altered by TmeA deletion (wild-type=0.106 fold/sec vs. ΔTmeA=0.092 fold/sec) but were reduced by over 10-fold following TarP deletion (ΔTarP=0.013 fold/sec) (Fig. 3E). Finally, we observed that ΔTmeA EBs achieved a significantly higher Fmn1 turnover rate than wild-type *Chlamydia* (Wild-type=0.041 fold/sec, ΔTmeA=0.070 fold/sec), while Fmn1 turnover was nearly negligible following TarP deletion (ΔTarP=0.009 fold/sec) (Fig. 3F). Thus, we conclude that the involvement of Fmn1 toward invasion-associated actin remodeling is TarP-specific, and that failure to recruit Fmn1 results in a substantial defect to both actin kinetics and *Chlamydia* internalization.

### Arp3 recruitment and turnover are dependent on TarP and TmeA, respectively

In addition to the presently established role of formins as enhancers of actin kinetics, it has also been established that the actin branching protein Arp2/3 is recruited by *Chlamydia* to drive actin recruitment (Lane et al., 2008). Indeed, a recent study suggested that TarP and TmeA signaling pathways form a functional collaboration that enhances recruitment of Arp2/3 and increases the rate of actin polymerization (Faris et al., 2020; Keb et al., 2021). As such, we monitored the recruitment of Arp3 fused to either GFP or mCherry at the entry sites of wild-type, *cis*-complemented, or effector deletion strains in order to assess the individual and combined contribution of the effectors on Arp2/3 recruitment (Fig. 4). We observed recruitment of Arp3 in all strains except for ΔTmeA/ΔTarP, indicating that Arp2/3 recruitment is contingent upon the presence of either TarP or TmeA (Fig. 4A,B). We noted that cis-TmeA EBs required an additional 20 seconds to recruit Arp3 at a comparable intensity to either wild-type or *cis*-TarP strains (Fig. 4B), which is consistent with the slight delay in peak actin intensity observed prior (Fig. 1B). Surprisingly, TmeA deletion did not substantially alter the intensity of Arp3 recruitment, whereas TarP deletion resulted in dim and transient Arp3 recruitment (Fig. 4B). In conjunction with the absence of Arp3 recruitment in the ΔTmeA/ΔTarP strain, these data suggest that both TarP and TmeA are independently capable of Arp3 recruitment. Nevertheless, the apparent differences between the Arp3 recruitment profiles of ΔTarP and ΔTmeA are inconsistent with a synergistic interaction between the effectors, instead indicating that Arp2/3 recruitment is primarily driven by TarP. However, we observed that Arp3 recruitment at the entry sites of ΔTmeA EBs was visibly more transient than Arp3 recruitment in either *cis*-TmeA or wild-type strains, indicating a potential role for TmeA in promoting stable recruitment of Arp2/3.

**Figure 4:**
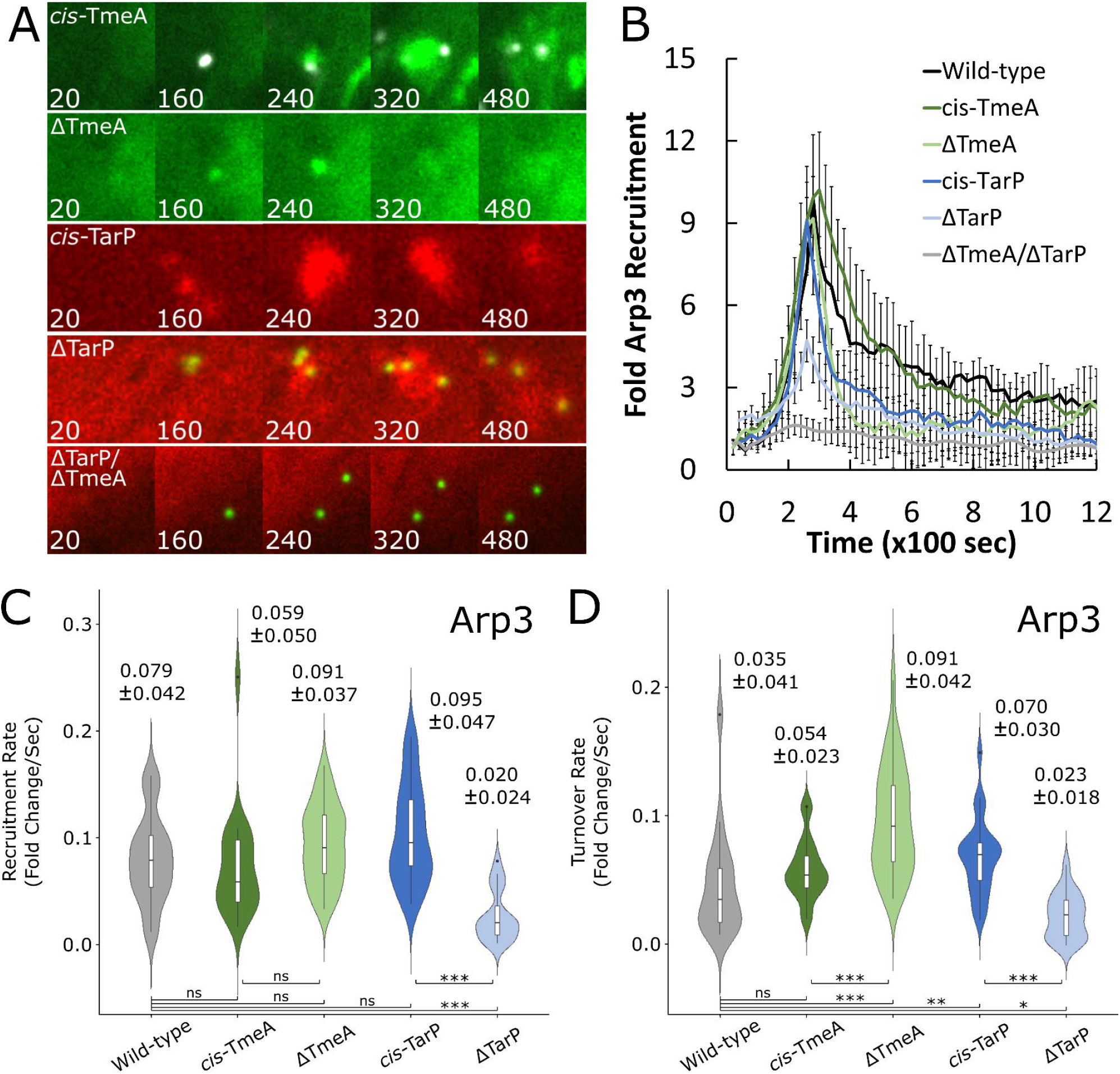
Arp3 recruitment and turnover are dependent on TarP and TmeA, respectively. (A,B) Cos7 cells were transfected with Arp3-GFP or Arp3-mCherry for 24 hours prior to infection with wild-type, single-knockout (ΔTarP, ΔTmeA), cis-complemented (*cis*-TarP, *cis*-TmeA), or double-knockout (ΔTmeA/ΔTarP) *Chlamydia* strains at MOI=20. Infection was monitored by live-cell confocal microscopy using a Nikon CSU-W1 spinning disk microscope, obtaining images every 20 seconds for 30 minutes to identify sites exhibiting Arp3 recruitment. (B) Fluorescence intensity of Arp3 recruitment was measured as described earlier (Fig. 1B) and plotted as mean fold recruitment for each timepoint +/-SEM compiled from a minimum N=12 recruitment events. (C,D) Kinetics of Arp3 recruitment and turnover were analyzed for each strain indicated above using the same methodology described in Fig. 1D,E. Violin plots contain a minimum N=12 individual rates, reporting the median rate +/-SD. Statistical significance was determined by Wilcoxon Rank-sum. All data are representative of at least 3 independent experiments, * P ≤0.05, ** P≤0.01, *** P ≤0.001.

In order to clarify the respective contributions of TarP and TmeA toward Arp2/3 recruitment, we monitored the kinetics of Arp3 recruitment and turnover for WT, ΔTarP, ΔTmeA, cis-TarP, cis-TmeA, and ΔTarP/ΔTmeA strains (Fig. 4C,D). We noted that the recruitment kinetics of Arp3 were largely similar in all strains except for ΔTarP and ΔTarP/ΔTmeA, the former recruiting Arp3 nearly 4-fold slower than wild-type or cis-complemented strains (Wild-type=0.079 fold/sec, ΔTarP=0.020 fold/sec) (Fig. 4C). Consistent with the delay in peak Arp3 intensity, *cis*-TmeA EBs possessed a slightly lower median Arp3 recruitment rate (*cis*-TmeA=0.059 fold/sec) than wild-type or *cis*-TarP EBs, although their overall kinetics are largely comparable. Finally, ΔTmeA EBs recruited Arp3 at a rate similar to wild-type or *cis*-TarP EBs (ΔTmeA= 0.091 fold/sec, *cis*-TarP= 0.095 fold/sec). Altogether, these data indicate that while TmeA is independently capable of recruiting Arp2/3, it does so at a far slower rate than strains which contain TarP. As such, we conclude that TarP is a key determinant of rapid Arp3 recruitment.

We found that the turnover rate of Arp3 was substantially higher at ΔTmeA entry sites (ΔTmeA=0.091 fold/sec) than wild-type (0.035 fold/sec) or cis-TmeA (0.054 fold/sec) (Fig. 4D). Furthermore, pairwise correlation analyses revealed that the Arp3 turnover rate of ΔTmeA EBs was positively correlated with both its corresponding recruitment rate and its recruitment intensity, together implying that rapid and intense Arp3 recruitment results in proportionally rapid Arp3 turnover (Fig. S9C). Collectively, this demonstrates that the reduced invasion efficiency determined for ΔTmeA is due to poor Arp2/3 retention at invasion sites resulting in attenuated actin kinetics. In other words, the inability of the ΔTmeA strain to maintain an optimal pool of activated Arp2/3 at the site of invasion could account for the observed two-fold reduction in the rate of actin recruitment. Meanwhile, TarP deletion resulted in a drastic attenuation of Arp3 turnover kinetics relative to wild-type or *cis*-TarP strains (*cis*-TarP=0.070 fold/sec, ΔTarP=0.023 fold/sec) (Fig. 4D). As with ΔTmeA, the significant reduction in actin recruitment by the ΔTarP mutant could be due to the drastic reduction in Arp2/3 recruitment, coupled with the loss of TarP-mediated recruitment of formins. In summary, these data indicate that TarP and TmeA signaling are mutually important for optimal Arp2/3 kinetics, each regulating distinct attributes of Arp2/3 recruitment. TarP signaling is associated with robust Arp2/3 recruitment, while TmeA signaling is required Arp2/3 retention at entry sites. Coupled with the attenuated actin kinetics and poor invasion efficiency of ΔTarP and ΔTmeA strains (Fig. 1), we conclude that efficient invasion is reliant upon the co-regulation of Arp2/3 kinetics by TarP and TmeA.

### Formin collaboration with the Arp2/3 complex is necessary for robust actin dynamics and efficient *Chlamydia* invasion

Having established that TarP signaling is necessary for robust recruitment of Arp2/3 and Fmn1, we opted to investigate the functional relationship between these two nucleators with regards to actin dynamics and consequent invasion. Findings from a previously published RNA interference screen in Drosophila S2 cells demonstrated that Arp2/3, formin, and Diaphanous are necessary for *Chlamydia* infection, but if and how they function together remained obscure. To assess more accurately the involvement of formin-mediated nucleation in actin dynamics and invasion efficiency, Arp2/3 complex participation must be excluded. As such, we compared the data obtained from SMIFH2-treated cells depicted earlier in the study (Fig. 3A-C) against Cos7 cells treated with the Arp2/3 inhibitor CK666 (100μM) or cells in which CK666 and SMIFH2 (100μM/10μM, respectively) were applied simultaneously. Cos7 cells were pre-treated with inhibitors for 60 min prior to inoculation with *C. trachomatis* serovar L2 EBs (MOI=50), quantifying invasion efficiency at 10 min post-infection. We normalized our data to the efficiency of invasion in absence of inhibitor in order to directly compare the effects of formin and/or Arp2/3 inhibition on invasion. Loss of activity of either formin or Arp2/3 yielded a comparable impediment to invasion, each reducing invasion efficiency by roughly 40%, while simultaneous inhibition of formin and Arp2/3 reduced invasion efficiency by 60% (Fig. 5A). The importance of formins and Arp2/3 toward efficient internalization are further underscored by the substantial 40-50% loss in invasion efficiency upon simultaneous knockdown of formins (Fig. 2A) or individual knockdown of Arp2 (Fig. S6). Together, these data suggest that while formin and Arp2/3 activity are independently important for invasion, their combined inactivation causes a more substantial inhibition of pathogen entry.

**Figure 5.**
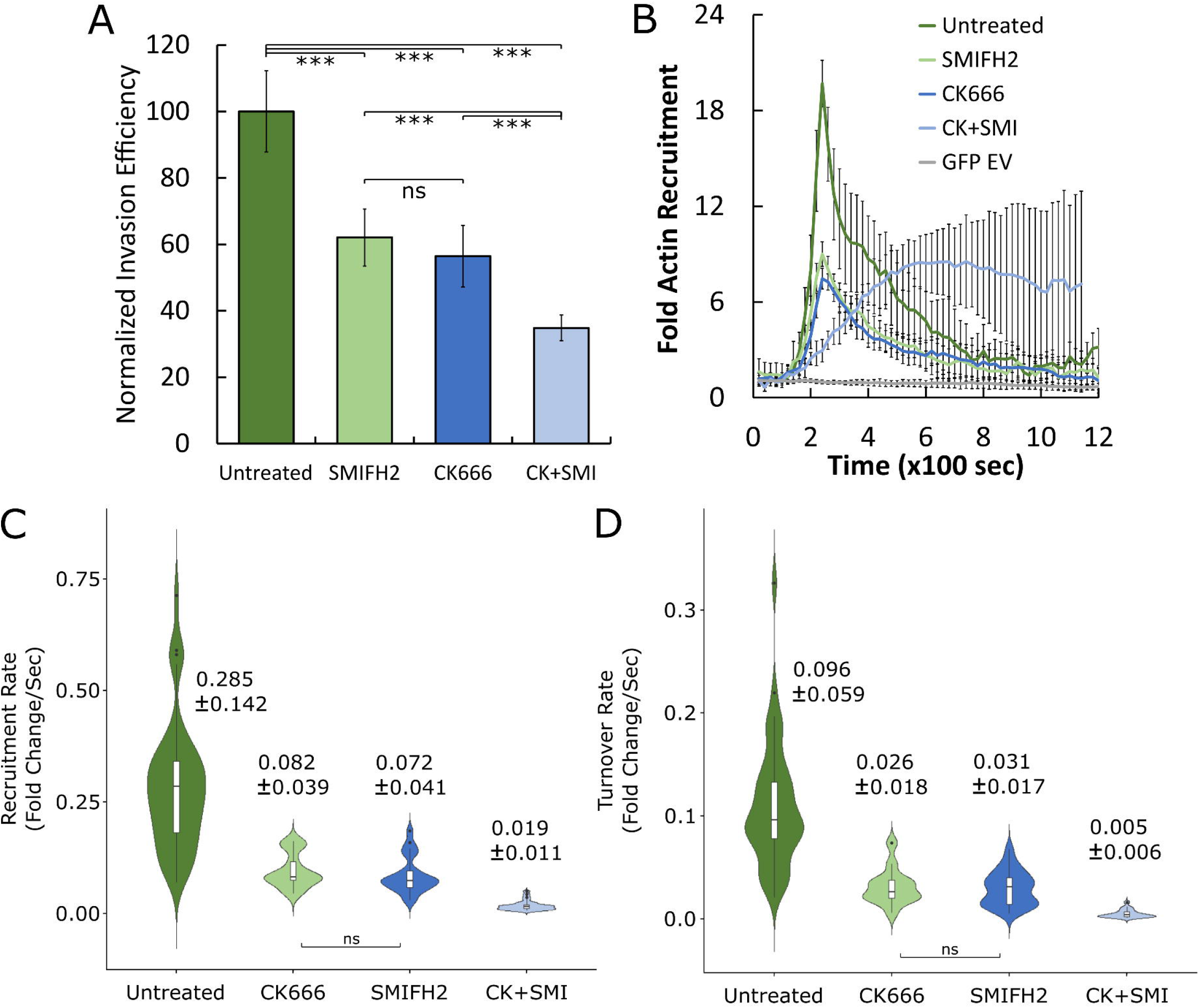
Formin and Arp2/3 activity are necessary for efficient invasion and optimal actin recruitment kinetics. (A) HeLa cells were mock-treated or pretreated with inhibitors against formin (10μM SMIFH2), Arp2/3 (100μM CK666) or both (CK+SMI) for 1hr, infected with *C. trachomatis* serovar L2 at MOI=50 and stained using the “in-and-out” method described earlier. Results were normalized against mean invasion efficiency of mock treated cells (Untreated mean = 49% invasion efficiency) and plotted as normalized mean +/- SEM. Data was collected from 20 fields, with each field containing an average of 108 Chlamydiae. Statistical significance was determined by T-test. (B) Cos7 cells were transfected with GFP-actin or a GFP empty vector plasmid for 24hrs prior to mock-treatment or pretreatment with 10μM SMIFH2, 100μM CK666, or both for 1hr. Transfected cells were infected with *Chlamydia* stained with CMTPX at MOI=20 and imaged by quantitative live-cell imaging, collecting images every 20 seconds for 30 minutes. Fluorescence intensity of GFP-actin recruitment was measured as described earlier (Fig. 1B) and plotted as mean fold recruitment for each timepoint +/-SEM compiled from a minimum N=16 recruitment events. (C, D) Kinetics of GFP-actin recruitment and turnover were analyzed for each condition using the same methodology described in Fig. 1D,E. Violin plots contain a minimum N=16 individual rates, reporting the median rate +/-SD. Statistical significance was determined by Wilcoxon Rank-sum. All data are representative of at least 3 independent experiments, * P≤ 0.05, ** P≤ 0.01, *** P ≤ 0.001.

We extended our earlier analysis of actin recruitment following the inhibition of formins (Fig. 3A-C) to include individual or co-inhibition of Arp2/3, quantifying the recruitment of GFP-actin in the presence or absence of CK666 and SMIFH2 via time-lapse imaging of live Cos7 cells (Fig. 5B). *Chlamydia* robustly recruited actin in the absence of inhibitor at an intensity of nearly 20-fold above background, which was reduced by over 10-fold following application of SMIFH2 or CK666. Interestingly, dual inhibition of formin and Arp2/3 yielded a form of actin recruitment that was substantially slower and much more persistent, often failing to turn over throughout the duration of the experiment (Video S3). In addition to exhibiting defects in actin recruitment intensity, we also found that the kinetics of actin recruitment and turnover were disrupted by inhibition of formin or Arp2/3 (Fig. 5C,D). Specifically, inhibition of formin or Arp2/3 comparably reduced the kinetics (fold/sec) of actin recruitment (Mock=0.285, SMIFH2=0.082, CK666=0.072) and turnover (Mock=0.096, SMIFH2=0.026, CK666=0.031), which was more severe upon co-inhibition of both nucleators. Altogether, these data indicate that *Chlamydia* manipulates both formin and Arp2/3 to achieve robust and highly localized actin recruitment, and that both nucleators must be functional to achieve efficient invasion. Additionally, via an unknown mechanism, actin turnover kinetics also required both Arp2/3 and formin nucleators to be functional.

The observation that formin and Arp2/3 individually contributes to actin kinetics during invasion raise the possibility that these nucleators act collaboratively, thereby enhancing the capacity of *Chlamydia* to generate actin-rich invasion structures. If nucleator collaboration does indeed play a role in invasion, we expect that both formin and Arp2/3 will be concurrently present within a single invasion event. We observed that GFP-Fmn1 was robustly recruited during invasion, but unexpectedly in an Arp2/3-dependent manner, in that CK666 treatment substantially reduced its presence at invasion sites (Fig. 6A). Likewise, GFP-Arp3, which has been validated as a marker for the Arp2/3 complex (Welch et al., 1997), was robustly recruited during invasion and also demonstrated reduced recruitment following inhibition of formin (Fig. 6B). Altogether, this indicates that recruitment of Fmn1 and Arp2/3 are reciprocally enhanced by their respective actin nucleating functions and subsequent actin polymerization.

**Figure 6:**
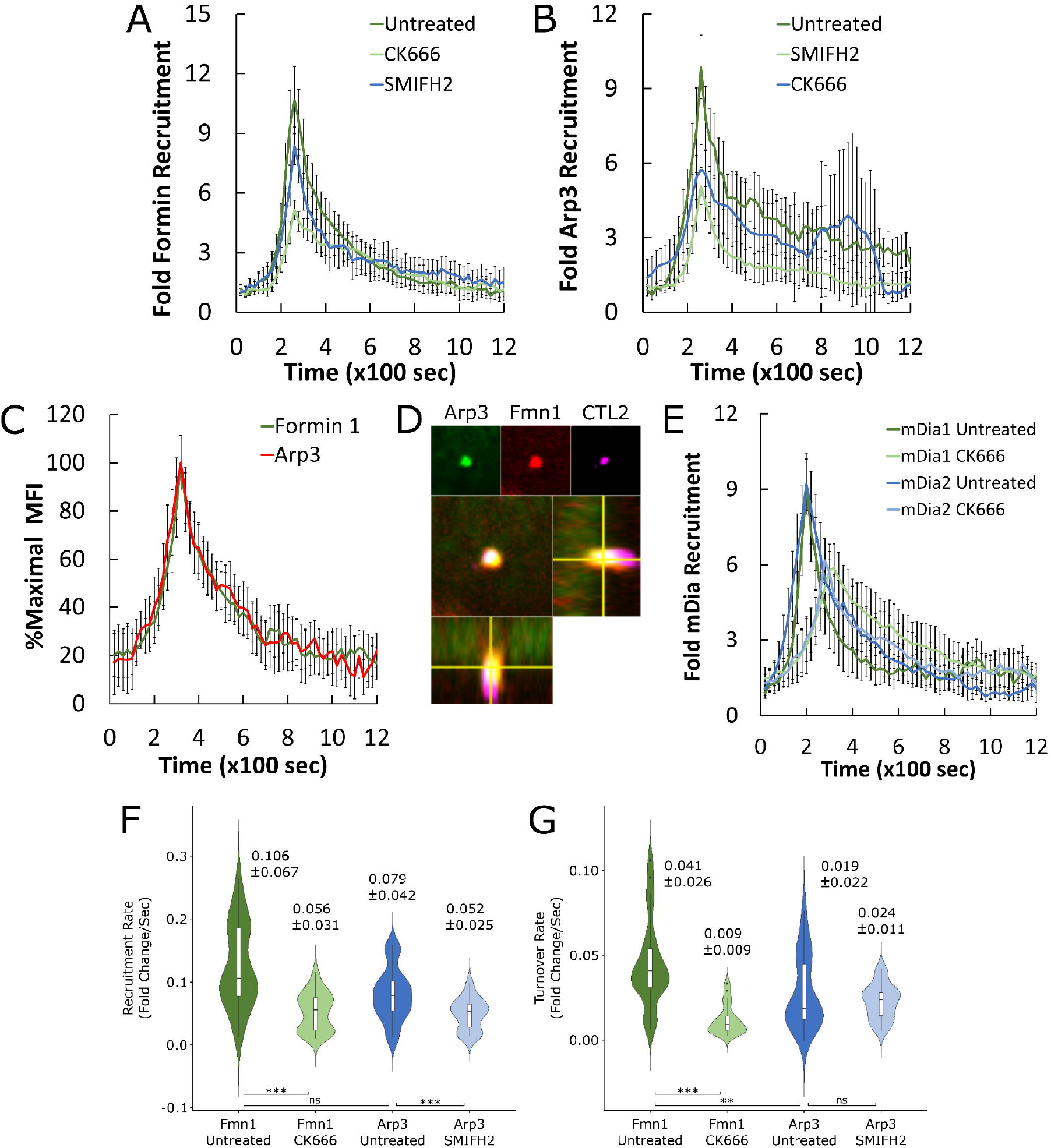
Formin 1 and Arp3 collaboration enhances recruitment kinetics at entry sites. (A,B) Cos7 cells were transfected with (A) GFP-formin 1 isoform 1B (GFP-Fmn1) or (B) GFP-Arp3 for 24 hours prior to mock-treatment or pretreatment with 10μM SMIFH2 or 100μM CK666 for 1 hour. Quantitative live-cell imaging of invading CMTPX-labeled *Chlamydia* (MOI=20) was performed as described in Fig. 1B. Briefly, images were acquired once every 20 seconds for 30 minutes and assembled into videos. MFI of recruitment events were obtained, and background fluorescence subtracted, before quantifying and plotting the fold recruitment values +/-SEM of Fmn1 or Arp3 for each timepoint. Data was compiled from a minimum N=16 recruitment events. (C) Cos7 cells were co-transfected with GFP-Fmn1 and mCherry-Arp3 for 24 hours prior to infection with unlabeled *C. trachomatis* (MOI=20) followed by quantitative live-cell imaging of invasion. Image acquisition and processing was performed as described above with a few modifications. Since CMTPX and mCherry-Arp3 fluorescence overlap, unlabeled *Chlamydia* was used, thus recruitment events are defined simply as regions containing elevated GFP-Fmn1 fluorescence compared to local background. MFI of both GFP-Fmn1 and mCherry-Arp3 were obtained from the recruitment ROI, and background was subtracted from both GFP and RFP channels independently. Maximal MFI was derived from both channels and used to normalize fluorescence as percent maximal MFI for each protein. Normalized MFI values for each timepoint were compiled from 33 events, plotting mean %Max MFI +/-SEM. (D) Cos7 cells were co-transfected with GFP-Arp3 and Fmn1-mCherry for 24 hours before synchronized infection with *C. trachomatis* (MOI=20). Invasion was halted at 20 minutes post-infection and *Chlamydia* stained via immunofluorescence. Cells were analyzed by confocal microscopy, obtaining Z-stacks of formin and Arp2/3 recruitment at sites of *Chlamydia* invasion. (E) Recruitment of mEmerald-mDia1 and mEmerald-mDia2 was monitored in the presence and absence of 100μM CK666 using the method described above for monitoring the recruitment of GFP-Fmn1 and GFP-Arp3. Data is displayed as mean fold recruitment for each timepoint +/-SEM compiled from a minimum N=22 recruitment events. (F,G) Kinetics of Fmn1 and Arp3 recruitment and turnover were analyzed in the presence and absence of 100μM CK666 or 10μM SMIFH2, respectively, using the same methodology described in Fig. 1D,E. Violin plots contain a minimum N=16 individual rates, reporting the median rate +/-SD. Statistical significance was determined by Wilcoxon Rank-sum. All data are representative of at least 3 independent experiments, * P ≤0.05, ** P≤0.01, *** P ≤ 0.001.

Next, we determined whether Fmn1 and Arp2/3 are co-recruited within individual invasion events, thus providing the necessary context for nucleator collaboration. To do this, we co-expressed GFP-Fmn1 and mCherry-Arp3 and measured the recruitment of both proteins during invasion by time-lapse imaging followed by fluorescence quantification (Fig. 6C). We observed that all events we monitored exhibited both Fmn1 and Arp3 accumulation, indicating that they are co-recruited by *Chlamydia*. To measure the timing of their arrival, we normalized the intensity of recruitment to the maximum fluorescence intensity of each protein at each timepoint, expecting that any delays in protein recruitment would be represented by a shift in their recruitment profile. In contrast, we noted that the recruitment profiles of both Fmn1 and Arp3 were remarkably similar and achieved maximal MFI at the same timepoint, indicating that both proteins arrived at the entry site within 20 seconds of each other, the temporal resolution of our imaging protocol. We confirmed these findings by collecting fixed-cell Z-stacks of formin and Arp2/3 recruitment at *Chlamydia* entry sites, noting that both nucleators were concurrently present at the base of invading elementary bodies (Fig. 6D). Given both the simultaneous recruitment of Fmn1 and Arp2/3 and the reciprocal dependence of their respective activities for optimal recruitment, we conclude that Fmn1 and Arp2/3 collaborate during invasion to promote actin recruitment. This observation is particularly novel, given that there are no other instances of the utilization of Fmn1 by bacterial pathogens within this collaborative context. Lastly, given the robust recruitment of Fmn1, we were interested in whether other formin species such as DRFs could also participate in nucleator collaboration. To this end, we found that mEmerald-mDia1 and mEmerald-mDia2 were also recruited during invasion, and that their recruitment was impaired by Arp2/3 inhibition (Fig. 6E, S7), suggesting that both DRF and non-DRF formins collaborate with Arp2/3 during invasion. Additionally, the capacity of mDia1 and mDia2 to participate in nucleator collaboration further supports our assertion that the multitude of formins recruited during invasion exhibit functional redundancy.

Using the same methodology to evaluate actin kinetics (Fig. 1D,E), we compared the recruitment and turnover profiles of Fmn1 and Arp2/3 to more closely evaluate how inhibition of each protein alters their recruitment and turnover dynamics during invasion. Compared to mock treatment, inhibition of Fmn1 decreased the rate of Arp3 recruitment by 34 percent (Mock=0.079 fold/sec, SMIFH2=0.052 fold/sec), while inhibition of Arp2/3 reduced the rate of Fmn1 recruitment by 47 percent (Mock=0.106 fold/sec, CK666=0.056 fold/sec) (Fig. 6F). Together, this indicates that collaboration between the activities of Fmn1 and Arp3 not only enhance the rate of actin recruitment (Fig. 5C), but also reciprocally enhance their respective recruitment rates. As such, we conclude that Arp2/3-mediated actin branching produces additional barbed-ends for Fmn1 binding; reciprocally, Fmn1 activity produces elongated actin filaments which allow efficient incorporation of Arp2/3-nucleated branches into actin networks generated by invading *Chlamydiae*. Fmn1-Arp2/3 collaboration was centered around TarP, but not TmeA, on the basis that Fmn1 recruitment was TmeA-independent (Fig. 3D) and that the ΔTmeA mutant (i.e. TarP-functional strain) exhibited similar cross-sensitivity toward nucleator inhibition as wild-type *Chlamydia* (Fig. S8). These data collectively implicate Fmn1 in efficient invasion via its collaboration with the Arp2/3 complex to mediate the hallmark robust actin dynamics. As an aside, the failure of TmeA to recruit Fmn1, despite the presumed availability of barbed ends in Arp2/3-nucleated actin network is interesting, and could indicate barbed end occupancy in TmeA-remodeled actin network, or that a TarP-specific cofactor of Fmn1 is required for its recruitment.

Shifting our focus onto turnover, we found that CK666 treatment reduced the turnover rate of Fmn1 by 4-fold compared to mock-treatment (Mock=0.041 fold/sec vs. CK666=0.009 fold/sec), while SMIFH2 treatment did not significantly alter the rate of Arp3 turnover (Mock=0.019 fold/sec vs. SMIFH2=0.024 fold/sec) (Fig. 6G). Furthermore, we found that the turnover rate of Fmn1 among individual EBs was moderately correlated with its corresponding peak intensity (r=0.402), which was not observed upon inhibition of Arp2/3 (r=0.074) (Fig. S9E). This discrepancy may be due to a relative lack of Fmn1 binding sites coupled with poor actin turnover following inhibition of Arp2/3 (Fig. 5D), such that the turnover rate of Fmn1 is dictated by binding site availability in inhibitor-treated cells. In contrast, rapid turnover of Fmn1 in mock-treated cells is potentially due to the relative abundance of Fmn1 binding sites coupled with rapid actin turnover, such that Fmn1 turnover is likely dictated by actin turnover dynamics. Meanwhile, we noted that SMIFH2 treatment did not alter the turnover rate of Arp2/3 (Fig. 6F), nor did it exert any observable effect on the regulation of Arp2/3 turnover as indicated by pairwise correlation analysis (Fig. S9E). Thus, we conclude that turnover of Fmn1 and Arp2/3 are governed by different mechanisms, wherein Arp2/3 turnover occurs via a process that is independent of Fmn1 activity.

## Discussion

In this study, we provide the first report of involvement of Fmn1, a member of formin nucleators, in *Chlamydia* invasion, indicating that its principal role is to collaborate with TarP-associated Arp2/3 recruitment to mediate the hallmark robust actin dynamics observed at sites of EB uptake. While the focus of this report is on actin recruitment, we also observed a correlation between reduced invasion efficiency and actin turnover, indicating that optimal actin remodeling, including disassembly of the actin network is an absolute requirement for efficient invasion. We provide ample evidence that TarP, which facilitates Fmn1-Arp2/3 collaboration is the major determinant of actin recruitment. While its role in invasion has been reported, here we were able to elucidate at the molecular level how it facilitates invasion. Its function as a signaling platform manifests as a facilitator of Fmn1-Arp2/3 cooperation. TarP appears to initiate Fmn1 and Arp2/3 recruitment, with accumulation facilitated by binding of nucleators to the growing actin network, which in turn contribute to further actin polymerization. Thus, a self-propagating model of recruitment of nucleators and actin explains the highly localized and robust actin dynamics during invasion. A crucial missing element is how this cycle is terminated to initiate turnover.

Intracellular pathogens often target the actin cytoskeleton during entry, leading to robust actin recruitment and subsequent formation of invasion-associated structures. Although *Chlamydia* has been demonstrated to use several entry mechanisms to achieve internalization (filopodial capture, clathrin-mediated endocytosis, macropinocytosis, membrane ruffling), we were unable to assign specific actin dynamic profile to each mechanism. Therefore, while our study impressed the importance of TarP and TmeA signaling toward optimal actin kinetics, we were unable to assign engulfment structures to a specific actin dynamics profile. Likewise, further study is required to determine the role of formin and Arp2/3 toward the assembly of each invasion-associated structure, and whether inhibition of either actin nucleator prevents the utilization of a given entry mechanism.

We observed that the presence of *Chlamydia* effectors TarP and TmeA was associated with rapid recruitment of actin, which was due in part to the collaboration between Fmn1 and Arp2/3. One explanation for the high rate of recruitment observed in our study is the establishment of a positive feedback loop, where actin branching by Arp2/3 and filament elongation by Fmn1 create additional nodes for Arp2/3 and Fmn1 to bind (Fig. 7). Iterations of this process would account for both the high recruitment rate of actin in addition to providing an explanation for why inhibition of Fmn1 and Arp2/3 comparably affect the recruitment rate of actin. However, this model does not immediately make apparent why actin recruitment is transient, nor does it provide explanation for the slow and persistent actin recruitment observed following inhibition of both formin and Arp2/3. Other host actin nucleators such as SPIRE, APC, or Cordon bleu have not yet been identified in siRNA screens of invasion-associated proteins, nor has any evidence of their involvement in *Chlamydia* invasion been given (Elwell et al., 2016). Although TarP has been shown via pyrene-actin polymerization assay to serve as an actin nucleator, it remains unknown whether it also performs this function in cell lines or *in vivo* (Jewett et al., 2006). However, based on pharmacological inhibition of formin members and the Arp2/3 complex and the drastic impairment of the kinetics of recruitment and turnover indicate that actin nucleation is predominantly driven by host nucleators rather than TarP itself.

**Figure 7:**
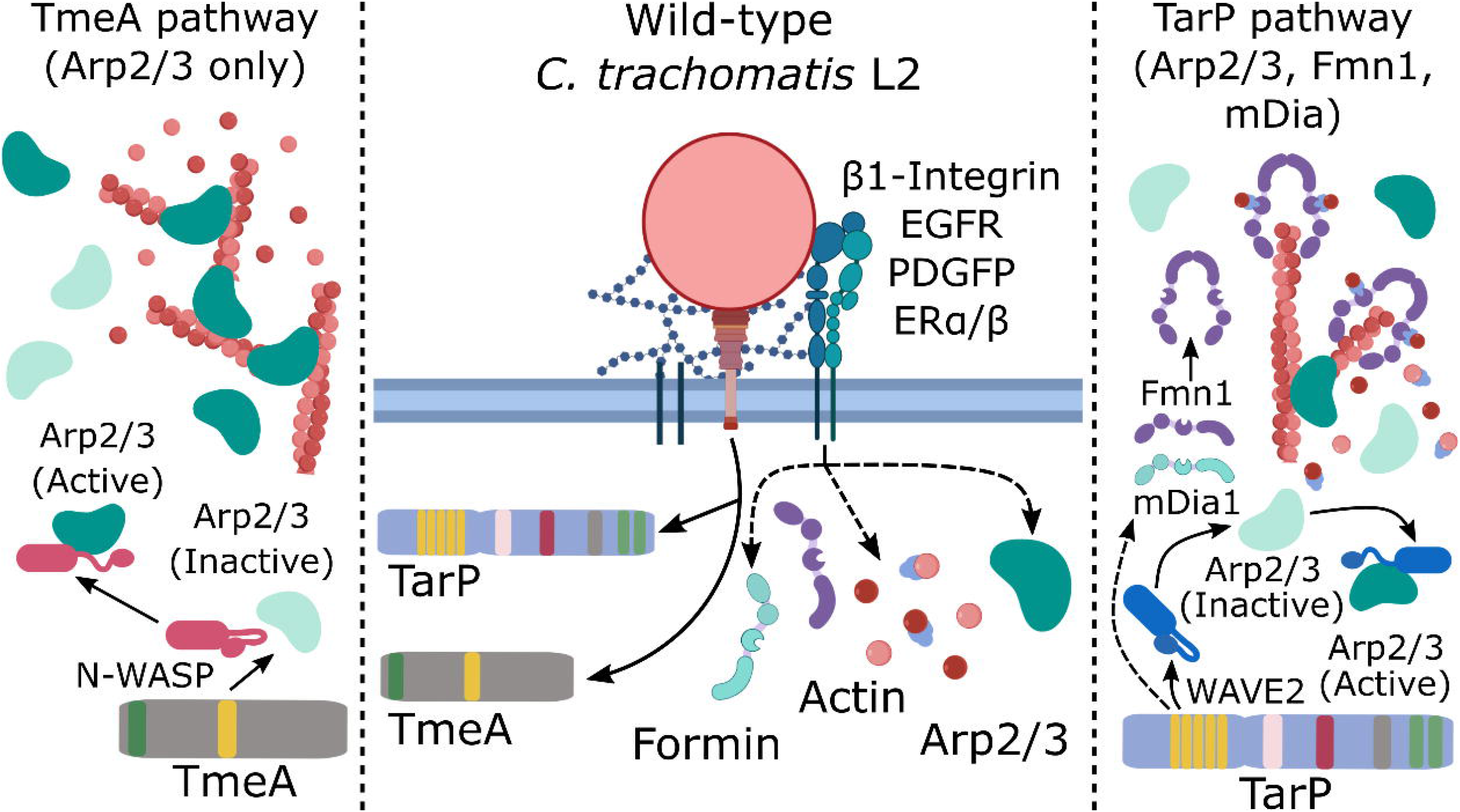
Proposed model for collaboration between formin and Arp2/3 and subsequent rapid actin recruitment and turnover. *C. trachomatis* engages and activates a multitude of host receptors whose activation is linked to the recruitment of formin, Arp2/3, and actin in non-invasion contexts. It is currently unknown whether receptor activation contributes to actin remodeling during invasion. Concurrent with receptor engagement, *Chlamydia* secretes the effectors TmeA and TarP via Type III secretion system; each effector activates Arp2/3 through N-WASP or WAVE2, respectively. Deletion of either TarP or TmeA attenuated the kinetics of Arp2/3 recruitment and turnover, which may reflect a failure to optimally activate and integrate Arp2/3 into actin networks. In addition to recruiting Arp2/3, TarP was also shown to be essential for the recruitment of Fmn1 and may also function to recruit mDia1 and/or mDia2 via signaling through Rac1. Collaboration between Arp2/3 and formin via their respective actin branching and elongating activities increases the rate of actin recruitment. Inhibition of Arp2/3 by CK666 or formin by SMIFH2 prevents this collaboration and constrains the actin network into either filamentous-only or branching-only networks.

An interesting observation is the apparent exclusive relationship of Fmn1-Arp2/3 collaboration with TarP. Why a TmeA-mediated Arp2/3 activation and ensuing actin nucleation and polymerization do not support Fmn1 localization to invasion sites is unclear. We speculate that the actin network formed by TmeA-N-WASP-Arp2/3 signaling is regulated by capping proteins that prevent Fmn1 access to barbed ends, or that additional factors are involved in TarP-dependent Fmn1 recruitment. The role of capping proteins have not been studied in detail in the context of *Chlamydia* invasion, but there are reports indicating that SIK2 and CAP1 capping protein regulators contribute to invasion *of Salmonella enterica* Typhimurium (Hahn et al., 2021; Misselwitz et al., 2011). In addition, the Drosophila homolog of the capping protein regulator CARMIL1 was identified in the siRNA screen by Elwell et al. to be necessary for *Chlamydia* infection of S2 cells (Elwell et al., 2008), thus raising the possibility that this protein regulates not only actin dynamics, but also the stable recruitment of Fmn1 within entry sites.

While TarP-mediated signaling accounts for the majority of actin recruitment, this is not reflected on invasion efficiency, where deletion of either TarP or TmeA resulted in similar levels of reduction. It is important to note that invasion is evaluated using an assay that relies on antibody accessibility. As such, even signaling pathways which yield only minor contributions to actin dynamics could result in substantial defects in *Chlamydia* internalization. One possibility is that TmeA-independent signaling is capable of robust initial recruitment of actin, but fails to provide the necessary signaling to achieve completion of engulfment. This observation is supported by our data indicating that the recruitment phase of TmeA-independent actin recruitment is comparable to wild-type *Chlamydia*, whereas actin turnover in the absence of TmeA is much faster, suggesting its involvement in a later stage of pathogen engulfment. Another possibility is that TarP and TmeA mediate different means of invasion, suggesting that their respective roles in achieving pathogen uptake entail some level of non-redundancy.

In addition to rapid actin recruitment, we also observed a two-phase actin turnover process starting with a brief fast turnover phase (100-180 sec) followed by a prolonged slow turnover phase (400-600 sec). The rate of actin turnover was reduced by deletion of either TmeA or TarP, or by inhibition of either formin or Arp2/3, collectively indicating that rapid actin turnover is an effector-driven process which is contingent upon the activity of actin nucleators. Although this process is relatively understudied in intracellular bacterial invasion, actin turnover is of clear importance for bacterial internalization. For instance, impairment or hyperactivation of the actin depolymerizing protein ADF-cofilin restricts internalization of *S. enterica*, while overexpression of the ADF-cofilin kinase LIMK reduced *L. monocytogenes* internalization (Bierne et al., 2001; Dai et al., 2004). In *C. burnetii* and *L. monocytogenes*, time-course experiments tracking actin recruitment also demonstrated a clear spike in actin recruitment followed by a two-phase rapid/residual turnover profile (Bierne et al., 2001; Meconi et al., 1998). Thus, the actin turnover profile observed in our study may simply reflect normal actin turnover dynamics, as opposed to being intrinsic to invading *Chlamydia.* Given the importance of host actin depolymerization factors like gelsolin, ADF-cofilin and their regulators for efficient internalization of other intracellular bacteria, it may be that such proteins are also responsible for actin turnover during *Chlamydia* invasion (Maldonado-Contreras et al., 2017; Moffatt et al., 2012; Pieters et al., 2013; Zheng et al., 2016). Indeed, an siRNA screen of *C. muridarum* invasion-associated proteins identified four actin depolymerization factors, ADF-cofilin, cyclase associated protein (CAP), Coronin, and β-thymosin, each of which reduced bacterial entry upon silencing (Fig. 7) (Elwell et al., 2008). Thus, one avenue for future study is to characterize the role of ADF-cofilin and other actin depolymerization factors in *Chlamydia* invasion, and detailed elucidation of their involvement would depend on investigations into actin dynamics.

Overall, this study highlights the remarkable complexity of the interaction between invading *Chlamydia* and the host cytoskeleton, identifying several novel interactions which culminate to support rapid actin kinetics and efficient pathogen engulfment. In play are at least two effectors (TarP and TmeA), signaling from accessory cell surface receptors, and the existing cross-regulatory machinery of actin nucleation, polymerization, and turnover (Fig. 7). Adding to this complexity is the various functional interactions between multiple formins/formin-like proteins with Arp2/3. Our data indicate that filamentous and branched actin nucleation along with polymerization constitute a fundamental interaction in which formins in general collaborate with Arp2/3 during invasion following their recruitment by TarP and TmeA.

## Materials and Methods

### Cell and Bacterial Culture

Green monkey kidney fibroblast-like (Cos7) cells and cervical adenocarcinoma epithelial (HeLa) cells were cultured at 37°C with 5% atmospheric CO2 in Dulbecco’s Modified Eagle Medium (DMEM; Gibco, Thermo Fisher Scientific, Waltham, MA, USA) supplemented with 10⍰μg/mL gentamicin, 2⍰mM L-glutamine, and 10% (v/v) filter-sterilized fetal bovine serum (FBS). HeLa and Cos7 cells were cultured for a maximum of 15 passages for all experiments. McCoy B mouse fibroblasts (originally from Dr. Harlan Caldwell, NIH/NIAID) were cultured under comparable conditions. *Chlamydia trachomatis* serovar L2 (434/Bu) was propagated in McCoy cells and EBs were purified using a Gastrografin density gradient as described previously (Caldwell et al., 1981).

### Reagents

SMIFH2 (Tocris, Minneapolis, MN, USA) and CK666 (Sigma-Aldrich, St. Louis, MO, USA) were diluted upon receipt to 100mM stock concentration in DMSO, S-nitro Blebbistatin (Cayman Chemical, Ann Arbor, MI, USA) and Y-27632 (Cayman) were diluted to 10mM stock concentration in DMSO. All inhibitors were dispensed into single use aliquots and stored at −20°C for no longer than 1 year after receipt. SMIFH2 was diluted to a working concentration of 10μM (1:10000), S-nitro Blebbistatin and Y-27632 were diluted to a working concentration of 10μM (1:1000), and CK666 was diluted to a working concentration of 100μM (1:1000), each using supplemented DMEM as diluent.

### Invasion Assay

Efficiency of *C. trachomatis* invasion in HeLa cells was performed as described previously (Carabeo et al., 2002). Briefly, HeLa cells were seeded in 24-well plates containing acid-etched glass coverslips and allowed to adhere overnight. Cells were pretreated with SMIFH2 (10μM) (Tocris, Minneapolis, MN, USA), CK666 (100μM) (Sigma-Aldrich, St. Louis, MO, USA), or both, for 1 hour prior to infection. Following inhibitor treatment, cells were infected with EBs derived from wild-type *C. trachomatis* L2 (434/Bu), *C. trachomatis* in which TarP, TmeA, or both were deleted by FRAEM (ΔTarP, ΔTmeA, ΔTmeA/ΔTarP), or *C. trachomatis* in which TarP or TmeA expression was restored by *cis*-complementation (*cis*-TarP, *cis*-TmeA) at MOI=50. EBs were allowed to attach to HeLa cells for 30 min at 4°C. Cells were rinsed with cold HBSS before temperature shift to 37°C by the addition of pre-warmed DMEM + 10% FBS/2⍰mM L-glutamine and incubated at 37°C for 10 min. Cells were then washed with cold HBSS containing 100 μg/mL heparin to remove any transiently adherent EBs before fixation in 4% paraformaldehyde at room temperature for 15 min. Fixed cells were labeled with a mouse monoclonal anti-*Chlamydia* LPS antibody (BioRad CF 6J12, Hercules, CA, USA), rinsed with 1x PBS, and fixed once more in 4% paraformaldehyde for 10 min. Next, cells were permeabilized using 0.1% (w/v) Triton X-100 for 10 minutes at room temperature, rinsed with HBSS and labeled with rabbit polyclonal anti-*Chlamydia trachomatis* antibody (Abcam ab252762, Cambridge, MA, USA). Cells were then rinsed in 1x PBS and labeled with Alexa Fluor 488 anti-mouse (ThermoFisher #A28175, Waltham, MA, USA) and Alexa Fluor 594 anti-rabbit (ThermoFisher #A-11012) IgG secondary antibodies. Coverslips were mounted and observed on either a Leica DM500 (Leica, Allendale, NJ, USA) or Zeiss Axio Observer (Zeiss, Dublin, CA, USA) epifluorescence microscopes. Percent invasion efficiency was quantified as total EBs (red) – extracellular EBs (green)/total EBs (red) x 100%.

### Quantitative live cell imaging of *Chlamydia* invasion

Cos7 cells were seeded onto Ibidi μ-Slide 8-well glass-bottomed chambers (Ibidi, Fitchburg, WI, USA) and allowed to adhere overnight prior to transfection. Cells were transfected with fluorescent proteins as indicated, using Lipofectamine 3000 (Thermo Fisher, Waltham, MA, USA) according to manufacturer directions. Transfection was allowed to proceed overnight before replacing media with fresh DMEM + 10% FBS/2⍰mM L-glutamine and allowing protein expression to continue for a total of 24 hours post-transfection. Transfection was verified on either a Leica SD6000 or Nikon CSU-W1 (Nikon, Melville, NY, USA) spinning disk confocal microscope prior to application of DMEM containing SMIFH2 (10μM), CK666 (100μM) or both. Wells were individually infected with CMTPX-labeled wild-type *C. trachomatis* L2 (434/Bu), unless otherwise indicated, at MOI=20 and promptly imaged using a 60x objective (NA 1.40) in a heated and humidified enclosure. Images were collected once every 20 seconds for 30 minutes, with focal plane maintained using an infrared auto-focusing system. Upon completion of the imaging protocol, the next well was infected and imaging repeated; mock-treated wells were imaged first to allow inhibitor treatment sufficient time to achieve inhibition. Images were compiled into videos using NIH ImageJ and analyzed to identify protein recruitment events. The mean fluorescence intensity (MFI) of such events was measured for each timepoint alongside the local background MFI of a concentric region immediately outside the recruitment event. Background MFI was subtracted from recruitment MFI for each timepoint and converted into fold recruitment by dividing the MFI of each timepoint against the baseline MFI, defined as the average MFI of five timepoints (100 sec) prior to the start of recruitment.

### Fixed cell confocal imaging

For Z-stacks of formin-1 and Arp3 colocalization, Cos7 cells were seeded onto Lab-Tek II 4-well glass bottomed chambers (Nunc, Rochester, NY, USA) and allowed to adhere overnight. Cells were co-transfected with Arp3-GFP and Fmn1-mCherry using Lipofecatmine 3000 according to manufacturer directions. Transfection was allowed to proceed overnight before replacing media with fresh DMEM + 10% FBS/2⍰mM L-glutamine and allowing protein expression to continue for a total of 24 hours post-transfection. Unstained density-gradient purified wild-type *C. trachomatis* elementary bodies were added in cold growth medium at MOI=20 and allowed to sediment for 30 minutes in a 4°c shaking incubator to synchronize infection. Invasion of elementary bodies was initiated by addition of prewarmed growth media followed by incubation at 37°C with 5% atmospheric CO2 for 20 minutes. Invasion was halted by rinsing cells with cold HBSS twice before adding 4% paraformaldehyde to fix cells. Next, cells were permeabilized using 0.1% (w/v) Triton X-100 for 10 minutes at room temperature, rinsed with HBSS and labeled with rabbit polyclonal anti-*Chlamydia trachomatis* antibody (Abcam ab252762, Cambridge, MA, USA). Cells were then rinsed in 1x PBS and labeled with Alexa Fluor 647 anti-rabbit (ThermoFisher #A-21245) IgG secondary antibody. Z-stacks were obtained using a Nikon CSU-W1 confocal microscope, imaging 0.3 micron slices of GFP (Arp3), RFP (Fmn1), and Cy5 (*Chlamydia*) fluorescence. Z-stacks were analyzed in ImageJ. For paxillin and vinculin staining, Cos7 cells were seeded onto acid-etched glass coverslips placed within each well of a 24-well plate and allowed to adhere overnight. Cells were treated with various concentrations of the myosin II inhibitor blebbistatin (5-20μM), ROCK inhibitor Y27632 (5-20 μM), or pan-formin inhibitor SMIFH2 (5-40μM), incubating cells for 1 hour. Cells were then fixed in 4% paraformaldehyde for 15 minutes, permeabilized in 0.1% (w/v) Triton X-100 for 10 minute, and probed with mouse monoclonal anti-vinculin (Abcam ab130007) and rabbit monoclonal anti-paxillin (Abcam ab32084) antibodies. Cells were then rinsed in 1x PBS and labeled with Alexa Fluor 488 anti-mouse (ThermoFisher #A28175) and Alexa Fluor 594 anti-rabbit (ThermoFisher #A-11012) IgG secondary antibodies. Imaging of focal adhesion staining was conducted using a Nikon CSU-W1 confocal microscope and images were assembled in ImageJ.

### RNA interference

To obtain lysates for Western blotting, or RNA for qRT-PCR, HeLa cells were seeded onto tissue culture treated 24-well plates and allowed to adhere overnight. Cells were transfected with 25-100 nM Dharmacon SmartPool siRNAs directed against ACTR2 (Dharmacon L-012076-02-0005, Lafayette, CO, USA), 100 nM Mission esiRNAs directed against FMN1 (Eupheria Biotech EHU089531, Dresden, Germany), DIAPH1/mDia1 (Eupheria Biotech EHU109471), DIAPH3/mDia2 (Eupheria Biotech, EHU090271), or 25-100 nM Trilencer-27 Universal scrambled negative control (Origene SR30004, Rockville, MD, USA) using Lipofectamine RNAiMAX reagent (Thermo Fisher) according to manufacturer directions. Incubation was allowed to proceed for 48 hours before lysing cells in either 2x Laemmli buffer for Western blotting, or in TRIzol (Thermo Fisher) for RNA purification. For invasion assays employing RNA interference, the transfection procedure was repeated as stated above for cells seeded onto acid-etched glass coverslips, followed by processing cells for “inside-out” staining as described earlier. For quantitative live-cell imaging following RNA interference, cells were seeded onto Ibidi μ-Slide 8-well glass-bottomed chambers and transfected using the method stated above, with some slight modifications. Specifically, 24 hours after initial transfection with siRNA or esiRNA, cells were transfected with GFP-actin using Lipofectamine 3000 reagent according to manufacturer directions and allowed to incubate for an additional 24 hours. Transfected cells were then imaged according to the protocol stated above for quantitative live-cell imaging.

### RNA isolation and quantitative RT-PCR

HeLa cells were seeded onto 24-well plates and allowed to adhere overnight before transfection with siRNA or esiRNA as detailed above. Cells were lysed using TRIzol reagent, incubating lysates for 5 minutes before transfer into a 1.5mL Eppendorf tube. Chloroform was added to tubes and shaken vigorously before allowing tubes to resettle for an additional 5 minutes. Lysates were centrifuged at 10000x RPM for 5 minutes before transferring the top aqueous layer into a fresh Eppendorf tube. Isopropanol was added to the aqueous layer, mixed gently, and incubated for 5 minutes prior to centrifugation at 14000x RPM for 20 minutes. Liquid was decanted from tubes before addition of 75% ethanol, followed by centrifugation at 9500x RPM for 5 minutes. Liquid was decanted from tubes and pellets containing purified RNA were allowed to air-dry before addition of Tris-EDTA buffer to resuspend RNA. cDNA was generated from purified RNA by PCR amplification using a SuperScript IV kit (Thermo Fisher) according to manufacturer directions. cDNA was diluted 1:10 in nuclease-free H_2_O and added to PowerUp SYBR Green Master Mix (Thermo Fisher) with specific qPCR primers diluted to 500 nM to generate a master mix. Each experimental sample was assayed in triplicate and were run on an Applied Biosystems QuantiStudio 3 Real Time PCR System using the following protocol: 1.) 50.0°C/2 min (x1), 2.) 95.0°C/10 min (x1), 3.) 95.0°C/15 sec (x40), 4.) 60.0°C/ 1 min (x1). Primers were subjected to dissociation curve analysis to ensure that a single product was generated.

### Western Blotting

Lysates generated as described above were resolved via SDS-PAGE in 10% polyacrylamide gels at 120 volts for 1.5 hours or until the dye front has begun to evacuate the bottom of the gel cassette. Gels were transferred onto 0.45μM pore size nitrocellulose in 1x Towbin buffer + 10% methanol at 90 mA for 16 hours. Western blots were blocked in 5% bovine serum albumin for 1 hour, briefly rinsed in Tris buffered saline + 0.1% Tween-20 (TBST) and incubated with appropriate primary antibody for 1 hour. Blots were then washed three times for 5 minutes in TBST and probed with appropriate HRP-conjugated secondary antibodies for 1 hour. Protein bands were resolved by chemiluminescence using Immobilon Western HRP Substrate (Millipore Sigma, St Louis, MO, USA). Western blotting of myosin II phosphorylation was conducted by densitometry analysis comparing the ratio of Phospho-Myosin Light Chain 2 (Thr18/Ser19) antibody signal (Cell Signaling Technologies #3674, Danvers, MA, USA) to total Myosin Light Chain 2 antibody signal (Cell Signaling Technologies #3672), normalized against GAPDH-HRP antibody signal (Abcam ab9385). Quantification of Arp2 expression via Western blotting was conducted by densitometry analysis of anti-Arp2 antibody (Abcam ab47654) compared to β-actin loading control (Abcam ab49900)

### Plasmids and DNA preparation

pEGFP-Actin-C1 (Heinzen et al., 1999) was provided by Dr Scott Grieshaber (University of Idaho). pEGFP-N1-ACTR3 (Arp3-GFP) (Welch et al., 1997) was obtained from Dr Matthew Welch (Addgene plasmid #8462). Arp3-pmCherryC1 (Taylor et al., 2011) was a gift from Christien Merrifield (Addgene plasmid #27682). mEmerald-mDia1-N-14 (Addgene plasmid #54157), mEmerald-mDia2-N-14 (Addgene plasmid #54159), and mRuby-LifeAct-7 (Addgene plasmid #54560) were gifted by Michael Davidson. pEGFPC2-FmnIso1b (Zhou et al., 2006) was a gift from Philip Leder (Addgene plasmid #19320). pmCherryC1-FmnIso1b was generated by a variant of Gibson assembly termed FastCloning (Li et al., 2011) and included removal of EGFP from pEGFPC2-FmnIso1b and insertion of mCherry derived from Arp3-pmCherryC1. Kanamycin-resistant transformants were selected and propagated for plasmid isolation prior to sequence verification using an mCherry-Fwd sequencing primer supplied by Eurofins Genomics. All plasmids were isolated using MiniPrep DNA isolation kits (Qiagen, Valencia, CA, USA) following a variant protocol for DNA isolation termed MiraPrep (Pronobis et al., 2016). Following plasmid isolation, the eluate was precipitated by addition of 3M sodium acetate (Invitrogen, Waltham, MA, USA) at 10% (v/v) of eluate volume followed by addition of 250% (v/v) absolute ethanol calculated after addition of sodium acetate. The mixture was incubated at 4°C overnight and centrifuged at 14,000×g for 15 minutes at 4°C. Supernatant was removed and 70% ethanol was added, followed by centrifugation at 14,000×g for 10 minutes at 4°C. Supernatant was removed once more, and precipitated DNA was resuspended in nuclease-free H_2_O. Primers for assembly of pmCherryC1-FmnIso1b are as follows: (pEGFPC2) Fwd: 5’ GGCGGAAGCGGAAGC<u>ATGGAAGGCACTCACTGCACC</u> 3’, Rev: 5’ GCCCTTGCTCACCAT<u>GGTGGCGACCGGTAGCG</u> 3’; (pmCherryC1) Fwd: 5’ CTACCGGTCGCCACC<u>ATGGTGAGCAAGGGCGAGG</u> 3’, Rev: 5’ GTGAGTGCCTTCCAT<u>GCTTCCGCTTCCGCCG</u> 3’.

### Graphs and statistical analysis

Violin plots were made using the ggplot2 base package (version 3.1.0) as a component of the Tidyverse package (https://cran.r-project.org/web/packages/tidyverse/index.html)in rStudio (version 4.0.3). Wilcoxon ranked-sum tests to determine statistical significance between violin plots and tabulated statistical analyses of violin plots were generated using base R statistics and the moments package (version 0.14, https://cran.r-project.org/web/packages/moments/index.html) in rStudio and assembled in Excel (Microsoft, Redmond, WA, USA). Linear regression analyses, recruitment plots, invasion assays, and all statistics associated with these data (pairwise T-test, correlation coefficient, SEM) were performed in Excel. All graphs were assembled using the free and open-source software GNU Image Manipulation Program (GIMP, https://www.gimp.org/) and Inkscape (https://inkscape.org/). Proposed model for collaboration (Fig. 6) was assembled using BioRender (https://app.biorender.com/).

## Acknowledgements

We thank the members of the Carabeo lab for extensive feedback regarding the design, direction, and analysis of the study. We thank Dr. Ken Fields (University of Kentucky College of Medicine) for the kind gift of ΔTarP, ΔTmeA, ΔTmeA/ΔTarP and TarP/TmeA cis-complemented strains. This study was supported by funding from the U.S. National Institutes of Health, National Institutes of Allergy and Infectious Disease grant R01 AI065545 (R.A.C.). M.D.R. was further supported by a fellowship from the Seattle Chapter of Achievement Rewards for College Scientists (ARCS). The contents and views expressed within this publication are the sole responsibility of the authors.

## Author Contributions

M.D.R. and R.A.C. designed the experiments and wrote the manuscript. M.D.R. performed the experiments, and M.D.R. and R.A.C. analyzed the data.

## Data Availability

Datasets regarding the raw data and analysis of invasion assays, fluorescence microscopy and associated kinetic analyses are available in Dryad (https://doi.org/10.5061/dryad.0k6djhb0g).

---

**Figure S1: Detailed image analysis throughput used to evaluate the intensity of protein recruitment.**

Quantitative live-cell imaging of fluorescent protein recruitment was utilized several times in various conditions throughout this study, necessitating a consistent throughput to facilitate direct comparison. (A) Videos of *Chlamydia* invasion were assembled following image acquisition in ImageJ and analyzed to identify protein recruitment events. Protein recruitment was defined by regions exhibiting high fluorescence intensity (Yellow) as compared to local background (Red). When possible, fluorescent *Chlamydia* (CMTPX-*Chlamydia*, GFP-*Chlamydia*, RFP-*Chlamydia*) were used to ensure recruitment was specific to an invading bacterium. Regions-of-interest (ROIs) specific to recruitment were selected to include the entirety of the recruitment event across all timepoints, and a concentric region immediately outside of this region was selected as a background ROI. (B) The mean fluorescence intensity (MFI) of both recruitment and background ROI was obtained for each timepoint. (C) Background MFI was subtracted from recruitment MFI to normalize for local variations in fluorescence intensity and for photobleaching during image acquisition. Fold recruitment was calculated by dividing the MFI of each timepoint by the average MFI of 5 timepoints (100 sec, yellow dots) prior to the start of recruitment, defined as the point in which recruitment velocity begins to increase. (D) Fold recruitment values were plotted against time for each recruitment event. Multiple events were aligned according to peak recruitment for each event before calculating the mean fold recruitment across all events.

**Figure S2: Endogenous SiR-stained actin recruits comparably to exogenously expressed GFP-actin.**

(A) Recruitment of GFP-actin at sites of CMTPX-*Chlamydia* entry (Fig. 3A) were represented as a time-course montage to depict actin recruitment dynamics. (B) Cos7 cell monolayers were treated with 1μM SiR-actin alongside the broad-spectrum efflux pump inhibitor Verapamil (10μM) for 30 minutes prior to infection with CMTPX-*Chlamydia* (MOI=20). Recruitment of SiR-actin at *Chlamydia* entry sites was monitored as described earlier (Fig. 3A) and represented as a time-course montage. (C) Fluorescence intensity of actin recruitment was measured as described earlier (Fig. 3A) and plotted as mean fold recruitment for each timepoint +/-SEM compiled from a minimum N=10 recruitment events.

**Figure S3: Entry and exit points for each phase determined by velocity of recruitment and turnover.**

Each event used to generate the protein recruitment plots shown throughout this study were divided into three phases, recruitment, fast turnover, and slow turnover, in order to study their kinetics. Entry and exit points for these phases was determined by plotting the first derivative value (i.e., velocity) of protein recruitment for each timepoint. Start of recruitment was defined as 20 seconds after velocity began to increase (t=160s, dark green), since this timepoint was more consistently associated with recruitment across events than the preceding timepoint (t=140s). End of recruitment was defined as the timepoint immediately prior to when velocity is at its minimum, or 20 seconds prior to Vmin (t=240s, light green). Start of fast turnover was defined as the timepoint associated with Vmin (t=260s, dark gray), and the end of fast turnover was defined as the point in which turnover ceased to decelerate (t=340s, light gray). Start of slow turnover was defined as the timepoint immediately following the end of fast turnover (t=360s, dark orange), and terminated at the timepoint prior to when velocity crosses the x-intercept (t=760s, light orange). This analysis was conducted for all events independently and was used as the basis for linear regression analysis to determine the rate (slope) of recruitment, fast turnover, and slow turnover across multiple proteins, strains, and inhibitor treatment conditions.

**Figure S4: SMIFH2 does not inhibit myosin superfamily proteins at relatively low concentrations.**

Cos7 cells were treated with (A) the nonmuscle myosin II inhibitor s-nitro blebbistatin, (B) the rho-associated protein kinase (ROCK) inhibitor Y27632, or (C) the pan-formin inhibitor SMIFH2 at various concentrations for 1 hour. Cells were probed with anti-vinculin and anti-paxillin antibodies and stained using fluorescent secondary antibodies to identify the integrity of focal adhesions following inhibitor treatment. (D) Cos7 cells were treated with blebbistatin, Y27632, or SMIFH2 at various concentrations for 1 hour prior to lysis in Laemmli buffer. Lysates were resolved by SDS-PAGE followed by Western blotting and probed with antibodies reactive against phospho-myosin light chain 2 (p-MLC2 Thr18/Ser19), total myosin light chain 2 (t-MLC2), and GAPDH. (E) p-MLC2 abundance was quantified for each sample by ratiometric analysis of p-MLC2 intensity versus total MLC2 intensity, with each condition normalized against mock treatment. The concentration of SMIFH2 utilized to inhibit formins throughout this study (10μM) did not prevent the formation of intact focal adhesions or restrict phosphorylation of MLC2.

**Figure S5: RNA interference of formin species attenuates invasion efficiency and causes phototoxicity.**

(A-C) HeLa cells were transfected with 100ng esiRNA to repress expression of Fmn1, mDia1, mDia2 either individually or collectively, using a pooled mixture of all three esiRNAs (3x esiRNA), alongside a scramble RNA control series. Expression levels of (A) Fmn1, (B) mDia1, or (C) mDia2 were quantified by qRT-PCR, normalized against scramble RNA control. (D) Cos7 cells were transfected with a pooled mixture of 100ng each Fmn1, mDia1, and mDia2 esiRNAs (3x esiRNA) for 24 hours, at which point cells were transfected with GFP-actin and incubated for an additional 24 hours. Transfected cells were infected with CMTPX-*Chlamydia* (MOI=20), monitoring invasion via live-cell confocal microscopy, collecting images every 20 seconds for 30 minutes. Quantification of actin recruitment was prevented due to cell death as a result of increased phototoxicity following esiRNA treatment. (E) Cos7 cells were transfected with esiRNA or scramble RNA control as described above, prior to infection with wild-type *C. trachomatis* elementary bodies (MOI=50), halting infection by application of 4% paraformaldehyde at 10 minutes post infection. Cells were stained to distinguish internalized elementary bodies using the method described earlier (Fig. 1C). Results were normalized against mean invasion efficiency of mock-treated *Chlamydia* and plotted as normalized mean +/-SEM. Data was collected from 15 fields, with each field containing an average of 72 *Chlamydiae*. Statistical significance was determined by T-test.

**Figure S6: RNA interference of Arp2 expression reduces invasion efficiency and actin recruitment.**

(A) HeLa cells were transfected with concentrations of Arp2 siRNA ranging from 25nM to 100nM for 48 or 72 hours, alongside 100nM scramble RNA control. Cells were lysed in 2x Laemmli buffer and resolved via Western blot, probing blots with antibodies against Arp2, in addition to β-actin as a loading control. (B) Band intensity of Arp2 was measured for each condition and normalized against the intensity of ß-actin, plotting Arp2 expression levels as percent expression relative to Arp2 expression in untreated cells. (C) HeLa cells were transfected with 25nm siRNA against Arp2 or 25nM scramble RNA for 48 hours prior to infection with wild-type *C. trachomatis* elementary bodies (MOI=50), halting infection by application of 4% paraformaldehyde at 10 minutes post infection. Cells were stained to distinguish internalized elementary bodies using the method described earlier (Fig. 1C). Results were normalized against mean invasion efficiency of mock-treated *Chlamydia* and plotted as normalized mean +/-SEM. Statistical significance was determined by T-test. (D) Cos7 cells were transfected with 25nm siRNA against Arp2 or 25nM scramble RNA for 24 hours, at which point cells were transfected with GFP-actin and incubated for an additional 24 hours. Transfected cells were infected with CMTPX-*Chlamydia* (MOI=20), monitoring invasion via live-cell confocal microscopy, collecting images every 20 seconds for 30 minutes. Fluorescence intensity of actin recruitment was measured as described earlier (Fig. 1B) and plotted as mean fold recruitment for each timepoint +/-SEM compiled from a minimum N=10 recruitment events.

**Figure S7: Kinetics of mDia1 and mDia2 recruitment and turnover are attenuated by Arp2/3 inhibition.**

All mDia1/2 recruitment events used to create the averaged plot shown previously (Fig. 2D) were individually divided into recruitment, fast turnover and slow turnover phases (Fig. S3). Individual rates of (A) recruitment and (B) fast turnover were plotted on a violin plot with inset boxplot, reporting the median rate +/-SD for each inhibitor treatment condition. Violin plots contain a minimum N=22 individual rates. Statistical significance was determined by Wilcoxon Rank-sum. Data are representative of at least 3 independent experiments, ** P ≤ 0.01, *** P ≤ 0.001.

**Figure S8: TmeA-independent actin recruitment is driven by formin and Arp2/3 activity.**

(A) Cos7 cells were transfected with GFP-actin for 24hrs prior to mock-treatment or pretreatment with 10μM SMIFH2, 100μM CK666, or both for 1hr. Transfected cells were infected with ΔTmeA *Chlamydia* EBs at MOI=20 and imaged by quantitative live-cell imaging, collecting images every 20 seconds for 30 minutes. Fluorescence intensity of GFP-actin recruitment was measured as described earlier (Fig. 1B) and plotted as mean fold recruitment for each timepoint +/-SEM compiled from a minimum N=16 recruitment events. (B, C) Kinetics of GFP-actin recruitment and turnover were analyzed for each condition using the same methodology described in Fig. 1D,E. Violin plots contain a minimum N=16 individual rates, reporting the median rate +/-SD. Statistical significance was determined by Wilcoxon Rank-sum. All data are representative of at least 3 independent experiments, * P≤ 0.05, ** P ≤ 0.01, *** P ≤ 0.001.

**Figure S9: Pairwise correlation analyses of protein recruitment, turnover, and intensity.**

Recruitment and kinetics data from (A) Fig. 1, (B) Fig. 3, (C) Fig. 4, (D) Fig.5, (E) Fig. 6, and (F) Fig. S7 were compiled and analyzed by pairwise correlation analysis to identify the extent of co-regulation between protein recruitment, turnover, and intensity for each protein and condition.

**Table S1: Proteins identified as TarP-interacting proteins via yeast two-hybrid screen.**

The pB27-TarP(1-1005)-LexA bait plasmid was used to screen a cDNA library consisting of randomly primed HeLa cell protein baits using a proprietary high-throughput yeast-two hybrid screening service (Hybrigenics ULTImate Y2H Screen) to identify host proteins which interact with TarP.

**Movie S1: Representative video of fluorescent protein recruitment during *Chlamydia* invasion**

Cos7 cells were transfected with plasmids expressing various GFP-fusion proteins for 24 hours prior to infection with CMTPX-stained *C. trachomatis* EBs (MOI=20). Bacterial adhesion and entry was monitored by live-cell imaging using a spinning disk confocal microscope (Nikon CSU-W1 or Leica SD6000 AF), obtaining images every 20 seconds for 30 minutes to identify sites exhibiting protein recruitment. Images were assembled into videos in ImageJ and annotated to highlight recruitment events. The representative video shown here depicts GFP-actin recruitment at sites where CMTPX-*Chlamydia* contacts the host cell (arrows). Similar videos were assembled for all proteins monitored throughout the study, each yielding recruitment events that exhibit tight localization around fluorescent *Chlamydia*, enabling straightforward quantification of recruitment intensity and duration (Figs. S1-3).

**Movie S2: GFP expressed via empty vector plasmid is not recruited during *Chlamydia* invasion.**

Cos7 cells were transfected with a GFP-empty vector plasmid for 24 hours prior to infection with CMTPX-stained *C. trachomatis* EBs (MOI=20). Infection was monitored via live-cell confocal microscopy as described earlier. Images were assembled into videos in ImageJ and annotated to highlight sites where CMTPX-*Chlamydia* contact the host cell (arrows). Absence of GFP-empty vector recruitment at sites of entry demonstrate that recruitment is specific to the host protein fused to GFP.

**Movie S3: Co-inhibition of formin and Arp2/3 yields abnormal actin recruitment kinetics around invading *Chlamydia***

Cos7 cells were transfected with GFP-actin 24hrs prior to pretreatment with 10μM SMIFH2 and 100μM CK666 for 1hr. Transfected cells were infected with unlabeled *Chlamydia* at MOI=20 and imaged by quantitative live-cell imaging, collecting images every 20 seconds for 30 minutes. Images were assembled into videos in ImageJ and annotated to highlight recruitment events (arrows).

